# New germline Cas9 promoters show improved performance for homing gene drive

**DOI:** 10.1101/2023.07.16.549205

**Authors:** Jie Du, Weizhe Chen, Xihua Jia, Xuejiao Xu, Emily Yang, Ruizhi Zhou, Yuqi Zhang, Matt Metzloff, Philipp W. Messer, Jackson Champer

## Abstract

Gene drive systems could be a viable strategy to prevent pathogen transmission or suppress vector populations by propagating drive alleles with super-Mendelian inheritance. CRISPR-based homing gene drives, perhaps the most powerful gene drive strategy, convert wild type alleles into drive alleles in heterozygotes with the help of Cas9 and gRNA. However, achieving successful outcomes with these drives often requires high performance. Specifically, it is desirable to identify Cas9 promoters that yield high drive conversion rates, minimize the formation rate of resistance alleles in both the germline and the early embryo, and limit somatic Cas9 expression. Thus far, high-performance promoters have only been discovered in *Anopheles* species. In *Drosophila*, the *nanos* promoter avoids leaky somatic expression, but at the cost of high embryo resistance from maternally deposited Cas9. To improve drive efficiency, we tested eleven *Drosophila melanogaster* germline promoters in several configurations. Some of the new promoters achieved higher drive conversion efficiency with minimal embryo resistance, but none could completely avoid somatic expression like *nanos*. However, such somatic expression often did not carry detectable fitness costs when the promoter-Cas9 elements supported a rescue homing drive targeting a haplolethal gene, suggesting somatic drive conversion. Based on our findings, we selected two Cas9 promoter lines for cage experiments with a 4-gRNA suppression drive. While one promoter exhibited substantial somatic effects, leading to a low drive equilibrium frequency, the other outperformed *nanos*, resulting in the successful suppression of the cage population. Overall, these novel Cas9 promoters hold potential advantages for homing drives in *Drosophila* species and may also possess valuable homologs in other organisms.

## Introduction

Gene drive is a promising method to control pest insect populations and reduce the spread of vector-borne diseases. Engineered gene drives can have higher inheritance rates than the normal 50% Mendelian expectation, allowing them to increase in frequency and eventually spread through a whole population^1^. Depending on the design goal, gene drives can be classified into two categories, modification and suppression. Modification drives could spread a desired cargo gene or another change into the target specie’s genome, while suppression drives are designed to reduce or eliminate the target species population for health, ecological, or economic purposes^2^.

There are many types of gene drives, but CRISPR homing gene drive is the most widely studied and perhaps the most powerful. In heterozygotes with a homing drive allele, the wild type allele can be converted into a drive allele by homology related repair (HDR). This process is called “drive conversion” or “homing.” Biased inheritance occurs when germline cells are converted from drive heterozygotes to homozygotes. Alternatively, the wild type allele could also be converted into a resistance allele by end joining repair, which often mutates the DNA’s sequence, preventing recognition by the drive’s guide RNA (gRNA)^34^. Ideally, Cas9 cleavage and HDR is confined to germline cells in early meiosis. However, drive conversion and resistance allele formation are not necessarily spatially restricted to germline cells, but can also occur in somatic cells if Cas9 is expressed in such cells^5^. Temporally, such activity can also occur in germline precursor cells^6^ and in zygotes or early embryos from parental Cas9 deposition. Likely because of the larger relative size of female gametes, only maternal Cas9 deposition appears to occur regularly, and this process only forms resistance alleles rather than supporting successful drive conversion^6–8^. An ideal promoter for Cas9 results in a high drive conversion rate, low resistance allele formation rate, and low level of somatic expression. Regardless of when they form, resistance alleles in drives with a specific target gene can be categorized as functional or nonfunctional, depending on whether they disrupt function of the target gene (either by frameshift mutation or other sufficient change in the protein’s amino acid sequence). When the drive has a higher fitness cost than functional resistance alleles, the drive allele frequency will be reduced over time. Fortunately, functional resistance can often be avoided by using multiplexed gRNAs^9^ and conserved target sites^10^. Nonfunctional resistance alleles usually cannot outcompete a drive but can reduce its overall efficiency (see below).

Successful construction of homing drives has been achieved in many species, including yeast^11, 12^, mice^13^, the fruit fly *Drosophila melanogaster*^14, 15^, and the mosquito species *Anopheles gambiae*^10^, *Anopheles stephensi*^16^, and *Aedes aegypti*^17^. Some homing endonuclease genes (HEGs) containing specific enzyme cut sites have been tested, such as I-PpoI in *Anopheles*, but CRISPR/Cas9 is more flexible because its target sequence is determined by gRNA(s) rather than the nuclease itself^18^. Cas9 based drive efficiency tends to be quite high in yeast and *Anopheles* mosquitoes, but lower in most designs for *Aedes*, flies, and especially mice. Homing gene drives can take many forms^2^. In the most basic form, they are unconfined to any target population and would spread widely, but variants such as split drive systems^19^ and daisy chains^20, 21^ that separate Cas9 and gRNA elements can make them self-limiting, being eliminated from the population after initially spreading under at least some parameter regimes. Confinement to target populations can also be achieved by targeting population-specific alleles^22^ or by using tethered systems, where a confined type of drive provides the Cas9 for homing drives^23, 24^.

Aside from these variants, homing drives can be configured for either modification or suppression. Modification drives usually contain a cargo gene (though other methods are possible), exemplified by a drive in *A. stephensi*, where antipathogen effector genes targeting malaria parasites were successfully expressed^8^. Use of the *vasa* promoter here caused high rates of embryo resistance, but this was mitigated in a *A. gambiae* homing drive that used the *nanos* promoter for Cas9^25^. In *A. aegypti*, the *exu* and *nup50* promoters for Cas9 did not show high efficiency^26^, and low drive conversion was also found with *sds3* and *bgcn*^27^. Despite working well in *Anopheles*, the *nanos* and *zpg* promoters also did not achieve drive inheritance rates above 75%^28^. However, when tested with a 4-gRNA construct, some genomic insertion sites for split Cas9 lines using the *shu* and *sds3* promoters showed high drive efficiency^17^. Somatic expression appeared to be moderate to high in all these *A. aegypti* lines. The most effective modification drives usually contain a recoded rescue element for an essential target gene, allowing removal of nonfunctional resistance alleles. When the target gene is haplosufficient (a single wild-type or recoded drive copy is enough for viability), nonfunctional resistance allele removal is slower, but effects from embryo resistance from maternal deposition and somatic Cas9 cleavage will be modest. This was demonstrated in *A. stephensi*^16^. Targeting of a haplolethal gene (where two functioning copies are required for viability) will allow immediate removal of nonfunctional resistance alleles, but embryo resistance can also remove drive alleles. In a *D. melanogaster* example, embryo resistance was low enough to allow drive success^15^, though this could potentially be an issue in other systems.

Suppression drives typically target essential but haplosufficient genes without providing a rescue. Such drives eventually form homozygotes, which are nonviable or sterile, thereby removing drive alleles from the population. If the drive frequency reaches a high enough level, this can lead to population suppression, but drives that lack sufficient genetic load will instead reach an equilibrium frequency with the population persisting (genetic load refers to the level of reduction in the reproductive potential of the population at this equilibrium). In *Anopheles gambiae*, three female fertility genes were selected for constructing gene drive systems^29^. By targeting female-specific genes, a drive could achieve higher suppressive power because drive alleles are only removed in sterile females that lack wild-type alleles, rather than in both sexes. However, aside from functional resistance, these drives suffered from high levels of embryo resistance from strong maternal deposition due to their use of the *vasa2* promoter for Cas9. Additionally, somatic Cas9 expression rendered female drive heterozygotes mostly sterile. Both of these factors reduce the genetic load of a suppression drive, though this reduction is large only when the drive does not have exceptionally high drive conversion. To reduce somatic expression and embryo resistance, *zpg*, *nanos*, and *exu* promoters were tested in *A. gambiae*, inspired by homology to known germline *Drosophila* genes^30^. The *exu* promoter showed low cut rates, but *zpg* and *nanos* had similar drive conversion to *vasa2* with much less embryo resistance and less somatic expression as well. Together with a conserved target site to avoid functional resistance, a homing suppression drive with the *zpg* promoter was able to eliminate an *Anopheles* cage population^10^.

In *Drosophila*, the oldest HEG-based homing gene drives tested used a wide variety of promoters (including *β-Tub85D*, *Mst87F*, *Hsp70Ab*, *vasa*, *Act5C-P*, *aly*, *bgcn*, *rcd-1r*, and *CG9576*) and 3’ UTRs for different nucleases^31–34^. None of these achieved high efficiency, though *rcd-1r*, *hsp70Ab*, and *Act5C-P* were able to promote some drive conversion. Cas9 based systems using *vasa* performed better, albeit with high rates of embryo resistance and somatic expression^6^. The *nanos* promoter had similar performance, without apparent somatic expression^6, 7^. The *rcd-1r* promoter was also tested at two target sites with similar performance, though only drive conversion was evaluated^35, 36^. Some of these promoters, together with *exu*, were evaluated in another recent study, but these Cas9 genes used a T2A fusion to EGFP, as well as the P10 terminator element, either of which may substantially change gene expression patterns. Some achieved high drive conversion efficiency based on the gRNA target site, but embryo resistance and somatic expression were not evaluated. Thus, despite being a model organism, Cas9 promoters in *D. melanogaster* have achieved less efficiency than *Anopheles* and perhaps even *Aedes*, as showcased by a suppression drive experiment that avoided functional resistance alleles with *nanos*-Cas9, but failed due to inadequate drive conversion efficiency, high embryo resistance, and high fitness costs^14^.

In this study, to improve *Drosophila melanogaster* homing gene drive efficiency, we constructed and tested eleven germline Cas9 promoters in different configurations. Some new promoters resulted in higher drive conversion rate and lower embryo resistance rate, but none was able to avoid somatic expression like the *nanos* promoter. Furthermore, two Cas9 promoters were selected for cage experiments with a 4-gRNA suppression drive, one of which had significantly better performance than *nanos*, resulting in the successful suppression of the cage population. These results demonstrate that these new Cas9 promoters could be useful in *Drosophila* homing gene drive systems.

## Methods

### Plasmid construction

For plasmid cloning, reagents for restriction digest, PCR, and Gibson assembly were obtained from New England Biolabs; oligonucleotides from BGI and Integrated DNA Technologies; 5-α competent *Escherichia coli* from TIANTEN and New England Biolabs; and the ZymoPure Midiprep kit from Zymo Research. Plasmid construction was confirmed by Sanger sequencing. We provide annotated sequences of the final insertion plasmids and target genomic regions in ApE format^37^ at GitHub (https://github.com/jchamper/ChamperLab/tree/main/Cas9-Promoters-Homing-Drive).

### Generation of transgenic lines

Embryo injections were conducted by Rainbow Transgenic Flies or Fungene. Donor plasmids (Table S1) were injected into *w*^1118^ flies (=500 ng/µL) together with a gRNA helper plasmid (100 ng/µL) and TTChsp70c9 (450 ng/µL), which was used as the source of Cas9 for transformation. To expand populations, injected individuals were first crossed with *w*^1118^ flies with four females and two males in each vial. Their offspring with EGFP or DsRed fluorescence in the eyes, which usually indicated successful insertion of the transgenic cassette, were then crossed for several generations to obtain homozygotes. Adults expressing slightly brighter eyes were more likely to be homozygous.

### Fly rearing and phenotypes

All flies were cultured with Cornell standard cornmeal medium or with a modified version (using 10 g agar instead of 8 g, addition of 5 g soy flour, and without the phosphoric acid) in a 25°C incubator with a 14/10-hour day/night cycle. Flies were anesthetized with CO_2_ and screened for fluorescence using NIGHTSEA adapters SFA-GR for DsRed and SFA-RB-GO for EGFP. Fluorescent proteins were driven by the 3xP3 promoter for expression and visualization in the white eyes of *w*^1118^ flies. DsRed was used as a marker to indicate the presence of the split drive allele or a synthetic target drive, and EGFP was used to indicate the presence of the Cas9 allele or served directly as the synthetic target. In split yellow drive systems, males usually only show natural color or yellow body color for both body and wings. However, females were considered as ‘mosaic’ if their body dorsal stripes or wing color were mixed yellow and natural. Each individual could also have one or both fluorescence colors indicating the presence of drive (DsRed) or Cas9/functional target (both EGFP).

### Cage study

For the cage study, flies were housed in 25×25×25 cm mesh enclosures. A line that was heterozygous for the split homing suppression drive allele^14^ and homozygous for the supporting Cas9 allele was generated by crossing drive males to individuals with the Cas9 line for several generations, selecting flies with brighter green fluorescence (which were likely to be Cas9 homozygotes) and then confirming that the line was homozygous for Cas9 by PCR.

Males from this line (heterozygous for the split homing suppression drive and homozygous for Cas9) were crossed to Cas9 homozygotes, and similarly aged Cas9 homozygotes were also crossed to Cas9 homozygotes males in separate vials for two days. All males were then removed, and females were then evenly mixed and allowed to lay eggs in eight food bottles for two days. Bottles were then placed in cages, and eleven days later, they were replaced in the cage with fresh food. Bottles were removed from the cages the following day (so that future larger generations only laid eggs for one day per generation), and the flies were frozen for later phenotyping for adult numbers and fluorescence. The egg-containing bottles were returned to the cage. This 12-day cycle with nonoverlapping generations was repeated for each generation.

Flies were occasionally given an extra day to develop if the bottles were due for replacement before approximately half of pupae had visibly eclosed (usually, most pupae would eclose after one day of egg laying followed by eleven days of development). When the population was observed to fall down to low levels near the end of successful cages, the flies were given fewer food bottles in which to lay eggs. The number was set to still keep substantially lower relative density compared to the normal equilibrium population, and this had the effect of increasing survival of larvae by reducing bacteria growth in bottles compared to the potential situation at near-zero density. This created a more robust population at lower population density, reducing the Allee effect.

### Phenotype data analysis

Data were pooled from different individual crosses in order to calculate drive inheritance, drive conversion, germline resistance, embryo resistance, and other parameters. However, this pooling approach does not take potential batch effects into account (each vial is considered to be a separate batch, usually with different parameters, but sometimes with the same parent for egg count data, see Supplemental Data Sets), which could bias rate and error estimates. To account for such batch effects, we conducted an alternate analysis as in previous studies^9, 14, 15, 24, 38^. Briefly, we fit a generalized linear mixed-effects model with a binomial distribution (maximum likelihood, Adaptive Gauss-Hermite Quadrature, nAGQ = 25). This allows for variance between batches, usually resulting in slightly different parameter estimates and increased standard error estimates. This analysis was performed with R (3.6.1) and supported by packages lme4 (1.1-21, https://cran.r-project.org/web/packages/lme4/index.html) and emmeans (1.4.2, https://cran.r-project.org/web/packages/emmeans/index.html). The code is available on Github (https://github.com/MesserLab/Binomial-Analysis). The alternate rate estimates and errors were similar to the pooled analysis (see Supplemental Data Sets).

### Genotyping

For genotyping, flies were frozen, and DNA was extracted by grinding flies from SNc9XSGr1 and SNc9XSGr2 lines separately in 200 µL DNAzol (Thermo Fisher) and an appropriate amount of 75% ethanol solution. The DNA was used as a template for PCR using Q5 Hot Start DNA Polymerase from New England Biolabs according to the manufacturer’s protocol. The region of interest containing the promoter and 5’ UTR fragment was amplified using DNA oligo primers AutoB_left_S_F and Cas9_S1_R. This would allow amplification of the DNA fragment with a 30 second PCR extension time. After DNA fragments were isolated by gel electrophoresis, sequences were obtained by Sanger sequencing and analyzed with ApE software^37^.

### Fitness cost inference framework

To quantify drive fitness costs, we modified our maximum likelihood inference framework^39^. Similar to a previous study^14^, we analyzed our homing suppression drive targeting female fertility. The maximum likelihood inference method is implemented in R (v. 4.0.3)^40^ and is available on GitHub (https://github.com/jchamper/ChamperLab/tree/main/Cas9-Promoters-Homing-Drive).

In this model, we make the simplifying assumption of a single gRNA at the drive allele site. Each female randomly selects a mate, and the number of offspring generated is reduced in drive/wild-type females if they have a fecundity fitness cost. No offspring are generated if females lack any wild-type allele. In the germline, wild-type alleles in drive/wild-type heterozygotes can potentially be converted to either drive or resistance alleles, which are then inherited by offspring. The genotypes of offspring can be altered if they have a drive-carrying mother and if any wild-type alleles are present. These alleles then are converted to resistance alleles at the embryo stage with a probability equal to the embryo resistance allele formation rate.

## Results

### New Cas9 regulatory element selection and construction

In this study, we constructed several *D. melanogaster* Cas9 elements and homing drives to reduce resistance allele formation. Drive conversion (homing) takes places in germline cells by homology-directed repair, while resistance alleles can be formed if end-joining repair instead mutates DNA at the gRNA target site (Figure 1). Resistance allele formation can also occur post-fertilization in the zygote or early embryo in the progeny of drive females due to maternal deposition of Cas9 and gRNA^6–8^. After embryo development, leaky somatic Cas9 expression (together with gRNA, which is usually expressed from ubiquitously active U6 promoters) could result in additional drive conversion or resistance allele formation. Because embryo resistance and somatic expression often occur in only a fraction of cells (due to delayed cleavage in the case of embryo resistance), an individual could have a mosaic genotype due to variable Cas9 cleavage and repair outcomes in different cells.

**Figure 1.**
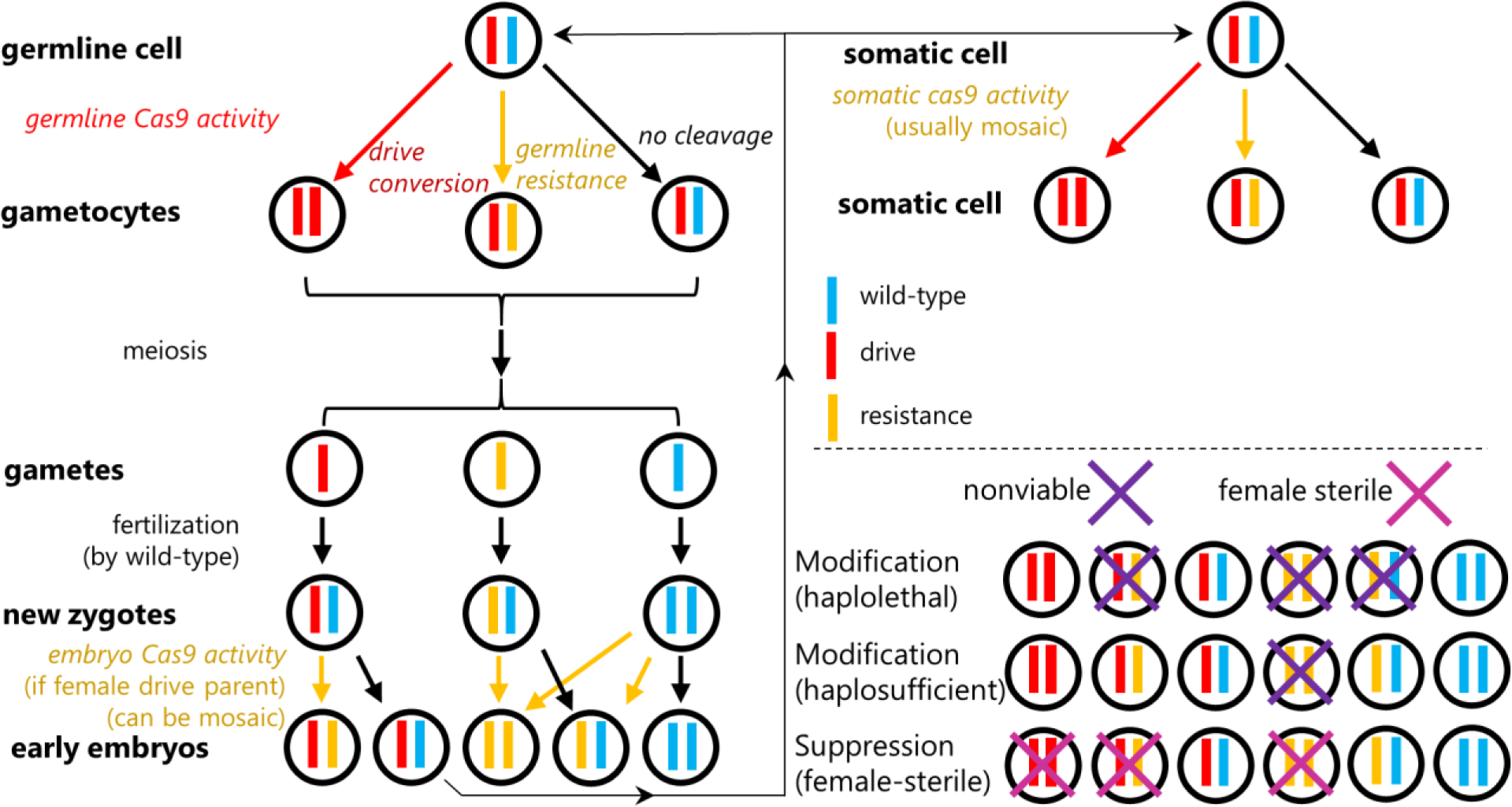
Cas9 activity in homing gene drive. Drive conversion occurs in germline cells of drive/wild-type heterozygotes. Cas9 cleavage can result in wild-type alleles being converted into drive alleles by homology-directed repair, but resistance alleles can also be formed by end-joining. After meiosis and fertilization, maternal deposition of Cas9 and gRNA can form additional resistance alleles in the zygote or early embryo, a process which can be mosaic. Somatic Cas9/gRNA expression later in development or in adults can also result in drive conversion or resistance allele formation, though this process appears to be independent of germline activity. Depending on the type of gene drive, certain individuals can be nonviable or sterile. In a rescue drive with a haplolethal gene target, any individuals with nonfunctional resistance allele will be nonviable. With a haplosufficient target, only individuals with two nonfunctional resistance alleles are nonviable. In female-sterile suppression drive, only females must have at least one wild-type allele (or functional resistance allele, not shown) to be fertile.

Resistance alleles can either preserve or disrupt the function of the target gene (referred to as functional and nonfunctional resistance alleles, respectively). Functional resistance alleles tend to be less common, because only one third of indel mutations from end-joining will preserve the reading frame, and many of the remaining alleles will be nonfunctional due to changes in the target protein’s amino acids. By using multiplexed gRNAs^6, 9^ and conserved target sites^10^, the fraction of functional resistance alleles can be greatly reduced.

In a haplolethal homing gene drive system designed for population modification, offspring that inherit any nonfunctional resistance gene will be nonviable (Figure 1) because a single working copy of a haplolethal gene is insufficient for viability. The drive allele contains a recoded “rescue” copy of the right portion of the target gene that preserves its function and cannot be cleaved by gRNAs. This drive system removes resistance alleles quickly, but it is vulnerable to embryo resistance and potentially somatic expression, which can result in drive alleles being removed in nonviable offspring. Another form of rescue drive targets a haplosufficient but essential gene. The only nonviable genotype for this is nonfunctional resistance allele homozygotes (Figure 1). This results in slower removal of nonfunctional resistance alleles, but avoids problems with somatic Cas9 expression and is only slowed, rather than stopped, by embryo resistance. A final type of drive is the suppression drive targeting a haplosufficient but essential female fertility gene without rescue. Here, males are unaffected, but females are only fertile if they have a wild-type allele. Drive/wild-type heterozygotes could have reduced fertility if there is somatic Cas9 expression. In general, functional resistance alleles have the same phenotype as wild-type alleles in all these drives, except that they would not be susceptible to somatic expression and cleavage if together with a drive allele (potentially reducing fitness costs in suppression drives and haplolethal rescue drives).

We constructed two types of drive systems. In our synthetic target drives, the homing drives are complete and target EGFP (Figure 2A) placed at “site C” on chromosome 2L^9^. The other system uses split Cas9 elements (Figure 2B and S1A) that are usually placed at “site B” on chromosome 2R^19^. These are then paired with one of three possible split drive elements for drive efficiency assessment. Each of these sites is downstream of two genes on either side to minimize fitness costs or other interference between genes. All drive elements have DsRed fluorescent markers, while Cas9 elements are marked with EGFP.

**Figure 2.**
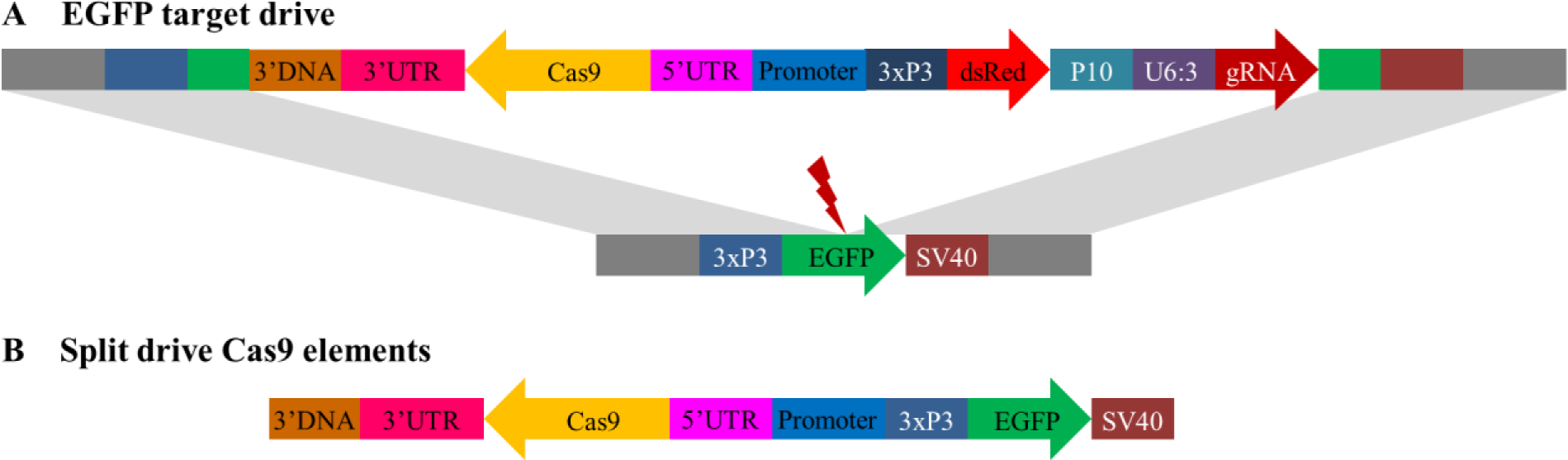
Schematic diagram of main constructs in the study. (**A**) The synthetic target drive is placed inside an EGFP gene at the gRNA target site. A DsRed fluorescence marker is regulated by the 3xP3 promoter for expression in the eyes together and a P10 3’ UTR element. A single gRNA driven by U6:3 promoter targets EGFP. Cas9 is driven by different compositions of promoter/5’ UTR and 3’ UTR (**B**) The split drive Cas9 elements all contain a EGFP fluorescent marker gene driven by the 3xP3 promoter and with a SV40 3’ UTR. Cas9 is driven by different compositions of promoter/5’ UTR and 3’ UTR.

Two of our drive systems are designed for easy visualization of nonfunctional resistance alleles, including the EGFP target drives and a split driving element targeting the X-linked *yellow* gene. Phenotypes for the EGFP drive are shown in Figure S1B, and functional resistance alleles are rare for this drive despite it having just one gRNA. For the X-linked drive targeting *yellow*^19^, null alleles have a recessive yellow body color phenotype (these can be drive or nonfunctional resistance alleles, see Figure S1C), and functional resistance allele represent approximately 10% of total resistance alleles^7, 19^. With our split Cas9 elements, we also tested a haplolethal drive targeting *RpL35A* with two gRNAs^15^ (Figure S1D) that have several nonviable genotypes, and also a suppression drive targeting the haplosufficient, female fertility *yellow-G* gene with four gRNAs^14^, which has several genotypes that are female sterile or reduced fertility (Figure S1E).

A list of all constructs used in the study can be found in Table S1, and Table S2 contains details of the sizes of our regulatory elements, including the promoter (defined here as DNA before the 5’ UTR), 5’ UTR, 3’ UTR, and included DNA downstream of the 3’ UTR. In general, the entire 3’ UTR were used, plus a small amount of additional DNA beyond the 3’ UTR in case this was important for transcription termination. For promoters, we used DNA that did not overlap with other genes (or an area immediately upstream of the 5’ UTR of other genes, which likely contained a core promoter of that gene), but in many cases, this would result in a very small promoter. In such cases, we often included 3’ UTRs of other genes, or even some of the 5’ UTR. For *nanos* and *vasa*, we used existing constructs as a basis^7^. Other promoters were selected for germline-restricted expression and low mRNA levels in the early embryo according to the Berkeley *Drosophila* Genome Project (https://insitu.fruitfly.org)^41^. *zpg* was selected due to its high efficiency in an *Anopheles* homing drive^10^, and *β2-tubulin* was selected because it is a known male-restricted germline promoter^42, 43^.

### Comparative drive performance at an EGFP target site

We designed and tested twelve Enhanced Green Fluorescence Protein (EGFP) target drives composed of different promoter and 3’ UTR elements in *D. melanogaster* (Figure 2A). These are similar to a drive described previously^9^ that used the *nanos* promoter/5’ UTR and 3’ UTR. Nearly all resistance alleles in this drive are nonfunctional, which disrupts EGFP and allows for determination of most drive performance parameters without sequencing (Figure S1B).

To determine drive performance, offspring from single drive/EGFP drive heterozygotes were phenotyped (Figure S2). In particular, female virgin drive/EGFP heterozygotes are crossed with males homozygous for EGFP. Drive/EGFP heterozygote males were crossed to EGFP homozygous or *w*^1118^ female virgin flies (Figure S2). Note that rather than reporting the standard parameter of drive conversion efficiency (the percentage of EGFP alleles converted into drive alleles in germline cells) and resistance allele formation rate, we report inheritance rates for compatibility with performance parameters for our haplolethal-targeting split drive below (where drive inheritance and conversion rates may not match normally due to potential drive-based differences in offspring viability).

Most drive systems showed 72-89% drive inheritance rates for males and 85-95% for females (significantly different from the Mendelian expectation, *P* < 0.0001 binomial exact test), with females having consistently slightly better performance (except for the one with the PEST sequence, see below) (Figure 3, Data Set S1). However, one drive system with the *β2-tubulin* promoter showed only Mendelian inheritance for both males and females. Even though *β2-tubulin* did not show any drive conversion, embryo resistance, or somatic activity, it still has some germline resistance formation in males. This is somewhat unexpected because the same promoter can support sex biasing from X-shredding^43^, which is thought to require relatively high cut rates to support multiple-cutting. For all other promoters, the total germline cut rate (drive conversion plus germline resistance allele formation) was usually 100% as measured in crosses between drive males and *w*^1118^ females. Drive inheritance rates for other constructs were generally similar. The drive with the *shu* promoter and 3’ UTR had the highest drive inheritance rate of almost 89% for males, and the drive with the *CG4415* promoter and *nanos* 3’ UTR had the highest drive inheritance rate in females of 95%. Only the drive with the *CG4415* promoter and 3’ UTR in males had a notably lower inheritance rate of 72%. Because these very different promoters showed similar germline performance despite likely having substantially varying expression levels, it is possible that Cas9 cut rates were highly saturated in the germline, perhaps due to use of a high activity gRNA.

**Figure 3.**
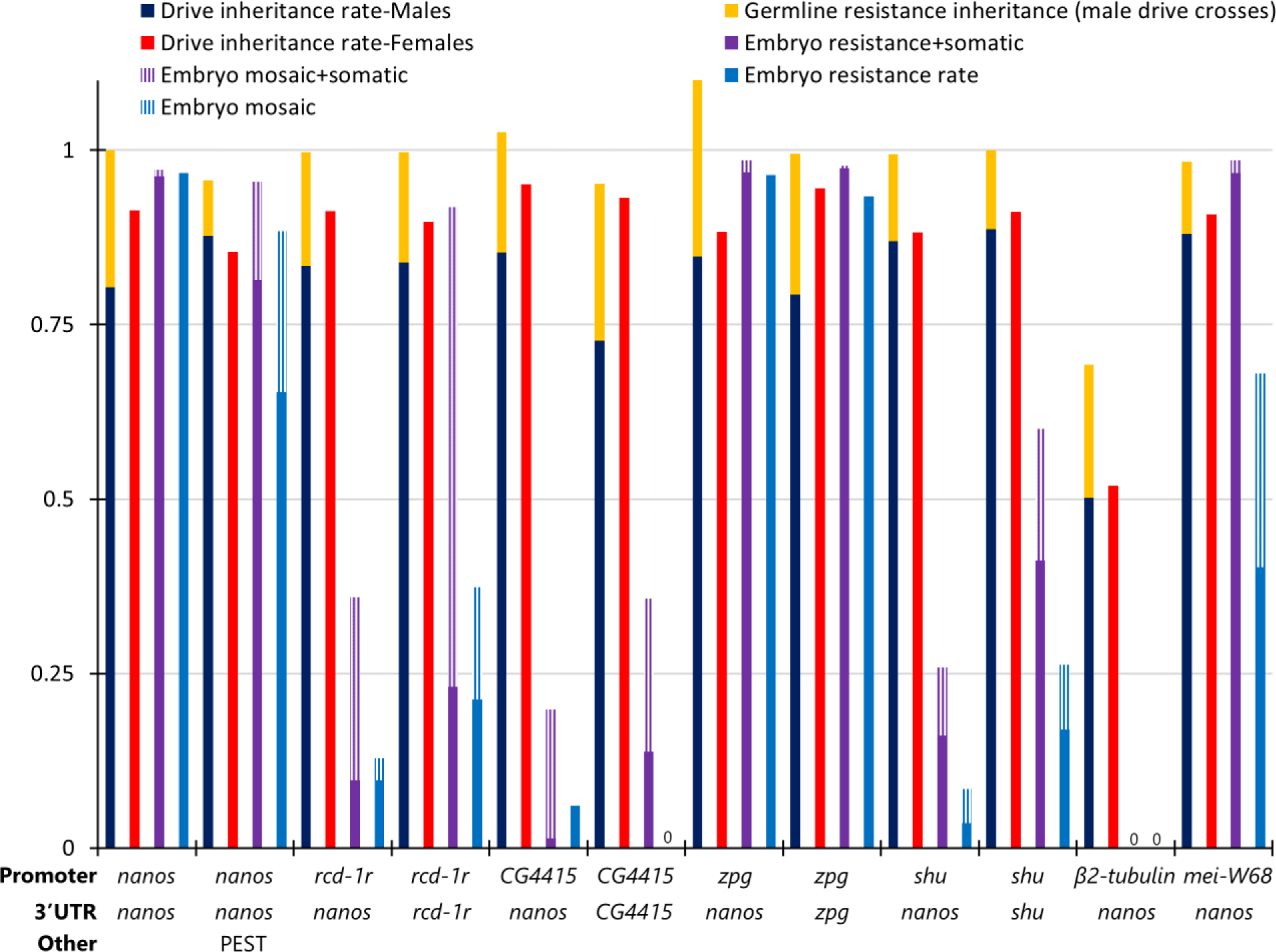
EGFP target site drive performance. The chart shows drive performance of twelve homing drive systems targeting EGFP on chromosome 2L differing by promoter/5’ UTR and 3’ UTR regulation of Cas9, or in one case, addition of a PEST sequence inside Cas9. Drive inheritance for males and females are measured in the progeny of drive/EGFP heterozygotes. Female germline resistance was not measured, and male germline resistance is measured from crosses with *w*^1118^ females (some male crosses were with EGFP homozygous females, and thus drive inheritance and germline inheritance can occasionally go above 1 because the pool of flies for the resistance rate was smaller). Drive inheritance rates above 0.5 shows that the ratio is above the normal Mendelian 50% rate. Female drive heterozygotes were always crossed with EGFP males. The fraction of offspring lacking EGFP phenotype (or with mosaic phenotype) and inheriting the drive is labeled as “Embryo resistance+somatic” because either maternally deposited Cas9 and gRNA or somatic expression in the eye could be responsible for lack of EGFP. “Embryo resistance rate” (and the corresponding mosaic rate) is similar, but reports the fraction of non-drive offspring lacking EGFP, which can only be caused by maternal deposition. 0 - no apparent phenotype in any offspring from embryo resistance or from combined embryo resistance and somatic expression. The leftmost drive data is from a previous study^9^.

Patterns in the embryo resistance rate (caused by maternal deposition of Cas9 and gRNA) in progeny of females varied more substantially between drive lines (Figure 3). However, this can only be directly measured in flies lacking a drive allele because somatic expression can also remove the EGFP phenotype or cause mosaicism. Except for the *nanos* promoter, which had low levels of somatic expression and mosaicism (and the small existing level was possibly due to proximity to the 3xP3 promoter), and *β2-tubulin*, which has low expression, all tested promoters showed moderate to high levels of somatic expression, resulting in mosaic drive/EGFP heterozygous parents. Mosaic phenotype was thus scored only for individuals that had at least 1/3 absence of EGFP in the surface of at least one eye, which was found to be sufficient to avoid scoring most individuals as mosaic (or with full embryo resistance alleles) when this was caused entirely due to somatic Cas9 expression, except for the *mei-W68* promoter, which had very high somatic expression. However, because germline cut rates were generally 100% in females, the embryo resistance allele formation rate could be measured directly in individuals that failed to inherit the drive (because they would still inherit a nonfunctional resistance allele from the mother, requiring additional cleavage only in the paternal EGFP allele). Some drive systems showed very high embryo resistances rates, such as those with the *nanos* and *zpg* promoters, and to a lesser extent the *mei-W68* promoter. This result for *zpg* is in contrast to its performance in *Anopheles gambiae*, where embryo and somatic expression were quite low^10^ (likely under 10% for embryo resistance^44^). The drives with *rcd-1r*, *shu*, and particularly the *CG4415* promoter showed much lower embryo resistance. For these, use of the *nanos* 3’ UTR tended to give slightly lower embryo resistance than the corresponding 3’ UTR of the promoters.

In the majority of flies inheriting the drive, lack of EGFP could be caused by either somatic expression or embryo resistance, but if the embryo resistance allele formation rate was high, then additional somatic expression would have little effect. Nevertheless, we saw notable increases in progeny that lacked EGFP phenotype or were mosaic in individuals inheriting drives with *rcd-1r*, *CG4415*, *shu*, and *mei-W68* promoters, which was less when the *nanos* 3’ UTR was used (Figure 3). Overall, the *rcd-1r*, *CG4415*, and *shu* promoters appeared to be promising combinations with the *nanos* 3’ UTR for high drive performance. These still had more somatic expression than the *nanos* promoter, but it was kept to a moderate level, and they had very low embryo resistance allele formation rates.

### Split drive performance at the *yellow* gene

Our EGFP target drives allowed an initial assessment of promoter performance in males and females, but they did have some disadvantages. First, they could only detect cutting activity in the eyes, but important somatic expression may be present in other tissues. Second, they made it difficult to distinguish between somatic expression and embryo resistance because most offspring inherited the drive, resulting in low sample sizes for calculation of embryo resistance. Third, they weren’t compatible with several newer split driving elements that were specialized for modification and suppression, representing drives closer to field applications. For our split drive systems, we designed and constructed several Cas9 elements, most at the same genomic locus. We first combined these with a split drive targeting *yellow* (Figure S1C), which tends to have somewhat lower embryo resistance than the EGFP drives^19^. It is also X-linked, allowing assessment of germline resistance inheritance from females (male offspring will only have one copy of *yellow* from their mother), and recessive knockout alleles cause a whole-body phenotype, allowing a different assessment of somatic expression. However, only drive performance in females can be tested.

Drive assessment was conducted by first crossing males homozygous for the Cas9 element to females that were homozygous for drive element (Figure S2). Then drive/Cas9 heterozygous female virgins were crossed with *w*^1118^ males. DsRed fluorescence for the drive element, EGFP fluorescence for the Cas9 element, and yellow body color phenotype were scored to assess drive performance (Figure 4, Data Set S2). In 17 of the 19 Cas9 elements with varying promoter and other factors, the drive inheritance rate mostly ranged 79% to 89% (significantly different from the Mendelian expectation, *P* < 0.0001 binomial exact test), but the *shu* promoter was only 71%, and the *CG17658* promoter had the lowest at 62%. The total apparent cut rate (drive conversion plus nonfunctional germline resistance allele formation) was usually very close to 100%, and the actual cut rate was likely 100% in many cases considering the relatively high functional resistance allele formation rate at this target site^7^ (such resistance alleles would appear as wild-type). However, the two drives with lower inheritance plus the *CG4415* promoter drive at “site C” (EGFP target drives were shown to have higher drive conversion when placed at this genomic site compared to our default “site B” locus^9, 19^) did not achieve complete germline cutting. The *nanos* promoter and 3’ UTR showed the highest drive inheritance of 88.7%.

**Figure 4.**
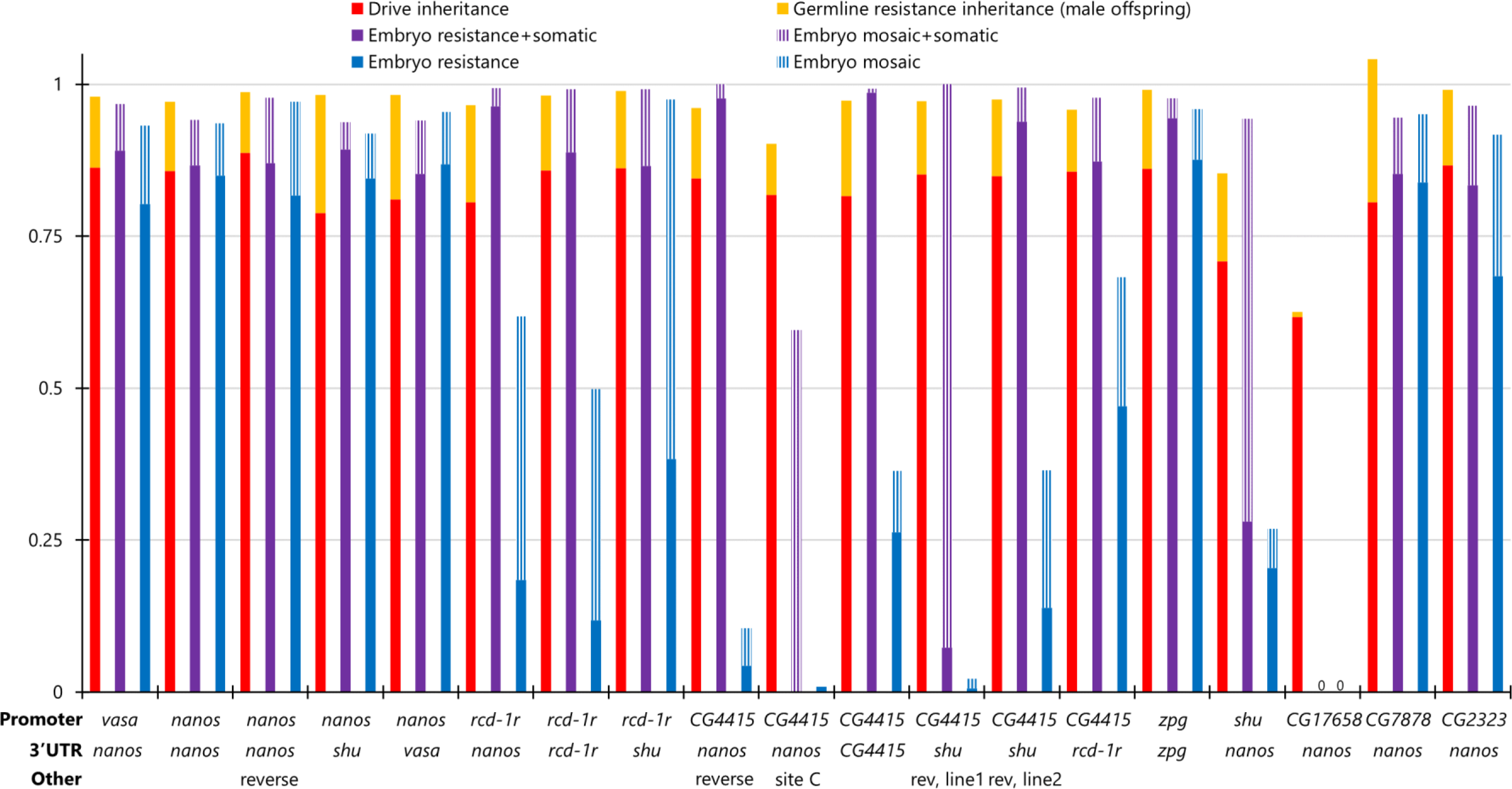
*yellow* target site drive performance. Females heterozygous for different Cas9 alleles on chromosome 2R (2L for “site C”) and heterozygous for a drive targeting the X-linked *yellow* gene were crossed with *w*^1118^ males. Their progeny were phenotyped for DsRed (drive), EGFP (Cas9), and yellow body color. The germline resistance inheritance rate shows the fraction of male progeny that had yellow body color but no drive allele (such flies could also form from embryo resistance allele formation). The fraction of offspring with yellow phenotype (or with mosaic phenotype), inheriting the drive, and also inheriting Cas9 is labeled as “Embryo resistance+somatic” because either maternally deposited Cas9/gRNA or somatic expression could be responsible for the yellow phenotype. “Embryo resistance rate” (and the corresponding mosaic rate) is similar, but reports the fraction of drive offspring lacking Cas9 that have the yellow phenotype, which can only be caused by maternal deposition. One Cas9 allele had different performance between lines, which are displayed as “line 1” and “line 2”. “Reverse/Rev” indicates that the orientation on one gene of the allele is reversed so that the Cas9 promoter and 3xP3 of EGFP promoter are not adjacent. 0 - no apparent phenotype in any offspring from embryo resistance or from combined embryo resistance and somatic expression.

Only three Cas9 promoter elements with high drive inheritance rates avoided high embryo resistance (Figure 4). These were *rcd-1r*, *CG4415*, and *shu*, though our test with *shu* showed less efficiency for drive inheritance. Of these, only *CG4415* with either the *nanos* or *shu* 3’ UTR (but not the *CG4415* 3’ UTR) had very low embryo resistance.

Somatic expression was more prevalent for the *yellow* split drive. In the initial cross, only drive heterozygous females with the *nanos* and *CG17658* promoters (the latter of which had very little activity in general), regardless of 3’ UTR, had no sign of any yellow mosaicism in any flies, which would be indicative of somatic Cas9 expression. For the *vasa*, *zpg*, *CG7878*, and *CG2323* promoters, somatic expression was moderate to high. For other lines, the level of somatic expression can be assessed by comparing female progeny with and without the Cas9 allele. Both can have embryo resistance alleles, but somatic Cas9 expression can only occur in progeny with a Cas9 allele. This allowed us to see moderate somatic expression in most remaining Cas9 lines based on the *rcd-1r*, *CG4415*, and *shu* promoters. In three lines based on the *nanos* and *CG4415* promoters, we reversed the orientation of the EGFP and Cas9 promoters to prevent the 3xP3 of EGFP potentially causing somatic expression of Cas9, which was a possible cause of the somatic expression seen with the *nanos* promoter in our EGFP target drives (we also saw fluorescent expression in the gonads of males and females with *nanos* adjacent to 3xP3, and in males with *rcd-1r* EGFP target drives, indicating the these promoters could affect each other). However, this was not necessary to avoid visible somatic expression with *nanos* for the *yellow*-targeting drive, and somatic expression remained in the *CG4415* lines (Figure 4). A more effective strategy involved placing Cas9 with the *CG4415* promoter at “site C,” which reduced somatic expression.

When assessing drive performance for these Cas9 elements, different lines were obtained, which usually showed the same performance and were thus combined in our analysis (a small number of lines showed no drive activity and were discarded). However, two sublines from with the *CG4415* promoter and *shu* 3’ UTR showed significantly different performance. The second line had notably higher embryo resistance and somatic expression (*P* < 0.0001 Fisher’s exact test). Genotyping detected no apparent difference in the insertion site, promoter/5’ UTR sequence, 3’ UTR sequence, or Cas9 itself, so it is unclear what caused the performance difference between these lines.

### Addition of a PEST domain for increased Cas9 degradation

One variant with the *nanos* promoter in our EGFP drives involved adding a PEST sequence to the C-terminus of Cas9. Such PEST sequences are known to increase the rate of protein degradation, and we hypothesized that this could reduce the level of effective maternal Cas9 deposition and thus reduce embryo resistance. Among progeny inheriting a drive allele, embryo resistance (somatic expression would not likely be a large factor in this *nanos* drive) was modestly reduced from 96% to 81%.

We thus decided to test several variants of split Cas9 elements with PEST sequences, some of which had reversed orientation between the Cas9 promoter and 3xP3 (Figure S2A). Unfortunately, these drastically reduced the effective drive inheritance rate (Figure S3, Data Set S3). While the embryo resistance rates and somatic expression levels were also reduced, the germline activity of this driving element perhaps was perhaps less highly saturated than the EGFP drives, resulting in the PEST addition causing a large reduction in germline cut rates, in addition to embryo activity. Even the strong *vasa* and *CG7878* promoters had drive inheritance rates of under 70%, and several drives were statistically indistinguishable from the Mendelian expectation (though all had at least some germline cleavage activity, and all except those with the *nanos* promoter had some noticeable somatic expression).

### Modification drive with haplolethal rescue performance

One common type of drive is the homing rescue drive, which allows modification drives to remove resistance alleles by targeting an essential gene. When the target is haplosufficient, drive dynamics are expected to be less strongly affected by promoter characteristics. When the target is haplolethal, any nonfunctional resistance alleles cause nonviability. Only individuals with drive and wild-type allele could survive. Thus, embryo resistance is more harmful, but resistance alleles can be removed much more quickly. The effect of somatic expression is less clear. If it tends to result in drive conversion in somatic cells, fitness effects may not be large. However, it is also possible that even mild somatic expression would form enough nonfunctional resistance alleles to induce severe fitness costs. We assessed these possibilities by combining eight of our split Cas9 lines with good performance with a previously constructed 2-gRNA haplolethal homing drive^15^, which also provides a good test for a drive with lower cut rates than our previous systems.

Drive homozygous females were crossed to Cas9 homozygous males, and the heterozygote progeny were then crossed to *w*^1118^ flies (Figure S2). In some cases, flies were allowed to lay eggs for 20-24 hours periods before being moved to each vial, and the eggs were counted to allow assessment of egg viability. Except for a Cas9 element driven by the *shu* promoter, all tested promoters showed high drive inheritance rates for males ranging from 87% to 93% (Figure 5, Data Set S4). However, only two Cas9 elements driven by the *nanos* and *CG7878* promoters had drive inheritance rates for females of over 70%. This is likely the result of reduced germline expression with these promoters, at least in females. For egg viability experiments, several controls were used of the same age and often in the same vials as the drive/Cas9 flies. The relative egg viability was assessed compared to these controls. Of the six promoters that underwent egg viability assessment (Figure 5), all had high egg viability in the progeny of males, ranging from 0.8 to 1.1 (with values above 1 likely representing stochastic fluctuations due to low samples sizes or minor differences in environmental conditions or fly health). In the progeny of males, nonviability is the result of germline resistance alleles, which likely occur at low frequency for this drive^15^, or fitness costs from somatic expression, which occur in the half of flies inheriting a Cas9 allele. Nonviability due to somatic expression would also be expected to reduce the frequency of Cas9 inheritance in the progeny of drive heterozygous males. This was not observed (Data Set S4), indicating that even when somatic expression is moderate (as in the *CG7878* promoter line), fitness costs are low. In the female lines, embryo resistance is also a factor. It is low for most of these lines, so the viability of the progeny of female drive individuals was also usually high. However, the *CG7878* promoter resulted in offspring viability lower than 0.1, most likely due to embryo resistance because the progeny of males were not affected.

**Figure 5.**
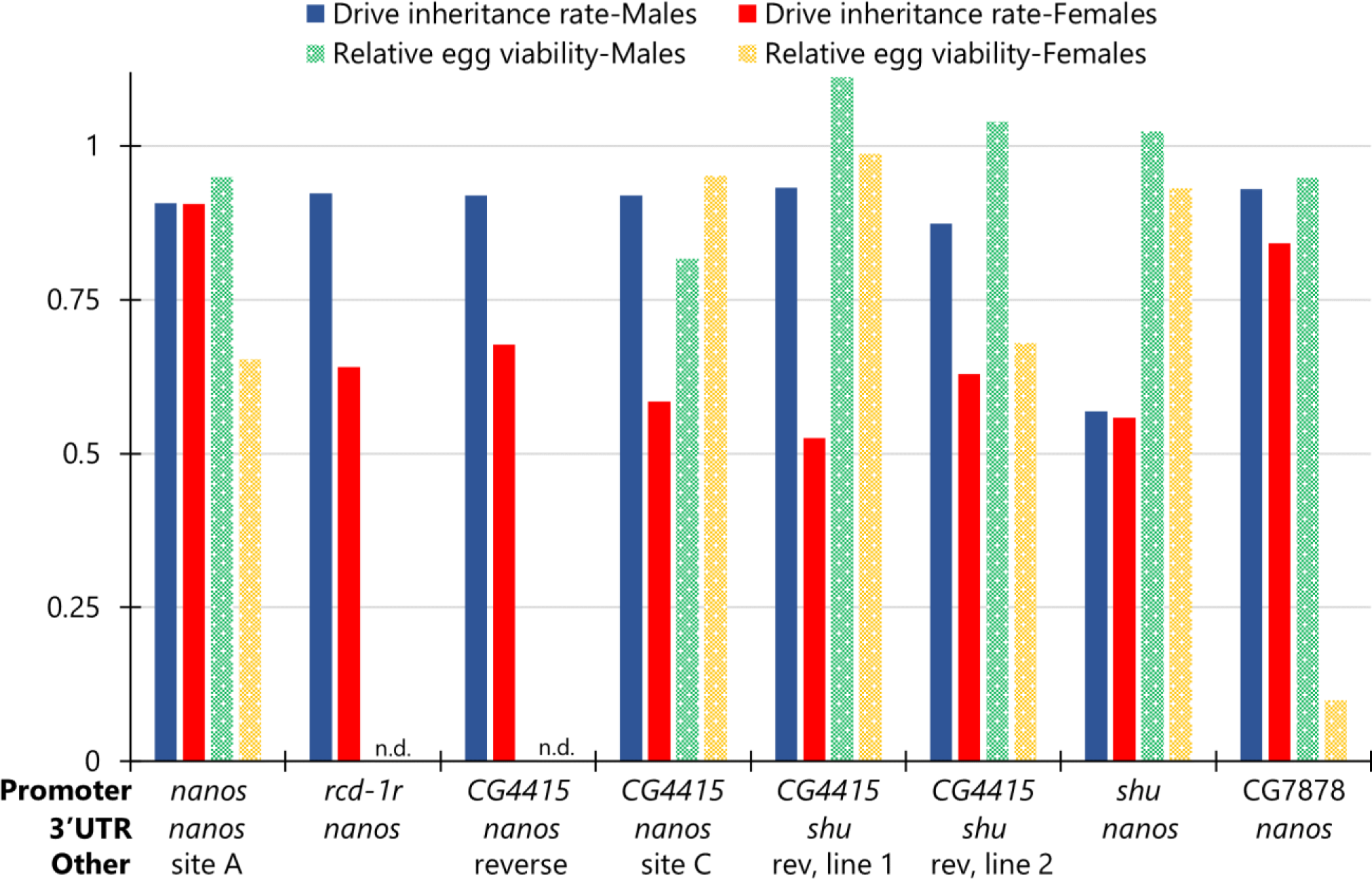
Drive inheritance and offspring viability of promoters with haplolethal homing rescue drive. Flies heterozygous for different Cas9 alleles on chromosome 2R (2L for “site C”) and a drive targeting the haplolethal *RpL35A* gene were crossed with *w*^1118^ flies. *RpL35A* is a haplolethal gene, so progeny with resistance alleles were nonviable, and high somatic expression could also potentially reduce viability of offspring that inherit both drive and Cas9. Progeny were phenotyped for DsRed (drive), and for several experiments, eggs were counted after one day of egg laying. Relative egg viability is the relative rates of egg survival compared to control experiments in which egg viability was measured for drive heterozygotes without Cas9, Cas9 heterozygotes, and crosses between *w*^1118^ flies. “Reverse/rev” indicates that the orientation on one gene of the allele is reversed so that the Cas9 promoter and 3xP3 of EGFP are not adjacent. n.d. - not determined. The leftmost drive data is from a previous study^15^.

### Suppression drive performance in individual crosses

Homing suppression drives targeting haplosufficient but essential female fertility genes have the potential to be the most powerful form of suppression drive with a high genetic load. However, if drive conversion is not very high, then embryo resistance and fitness costs in heterozygous females can reduce the suppressive power of the drive because the drive is not able to reach a high equilibrium frequency. This was the case with our 4-gRNA drive targeting *yellow-G*^14^, an egg shell protein that is critical for egg development. While fitness costs due to somatic Cas9 expression can have a severe effect (unlike in modification drives, drive conversion in somatic cells would also reduce fitness), this drive suffered fitness costs even with the *nanos* promoter, indicating that disruption of at least this specific target gene in the germline also reduces fertility. None of our new promoters had less somatic expression than *nanos*, but they often had less germline expression, potentially reducing fitness costs, and several have lower embryo resistance.

With a similar experimental setup to our investigation of the haplolethal split homing drive (Figure S2), we found that drive inheritance from males was also consistently higher than from females (Figure 6, Data Set S5), though differences were smaller than for the drive targeting *RpL35A*. Except for the *shu* promoter, which had worse performance, all other promoters had high drive inheritance rates for males ranging from 84% to 95%. In progeny of females, drive inheritance ranged from 67% to 87%. Eggs from male parents retained high viability (between 0.91 and 1.04). However, most females had lower egg viability (Figure 6). While the *nanos* promoter showed a small reduction, others had more substantial reductions, and somatic expression from the *shu* and *CG7878* promoters resulted in no eggs being viable. Only the *CG4415* promoter placed at chromosome 2L retained high egg viability. Though it has more somatic expression than *nanos*, this increased fitness may come from less germline expression, or perhaps a different spatial or temporal pattern of expression in the germline or ovaries in general. However, somatic expression remains important for fitness with this drive, as evidenced by the *shu* promoter, in which no eggs were viable despite low germline cut rates (at least in males, with a similar pattern for females in other drives - see Figures 4-5).

**Figure 6.**
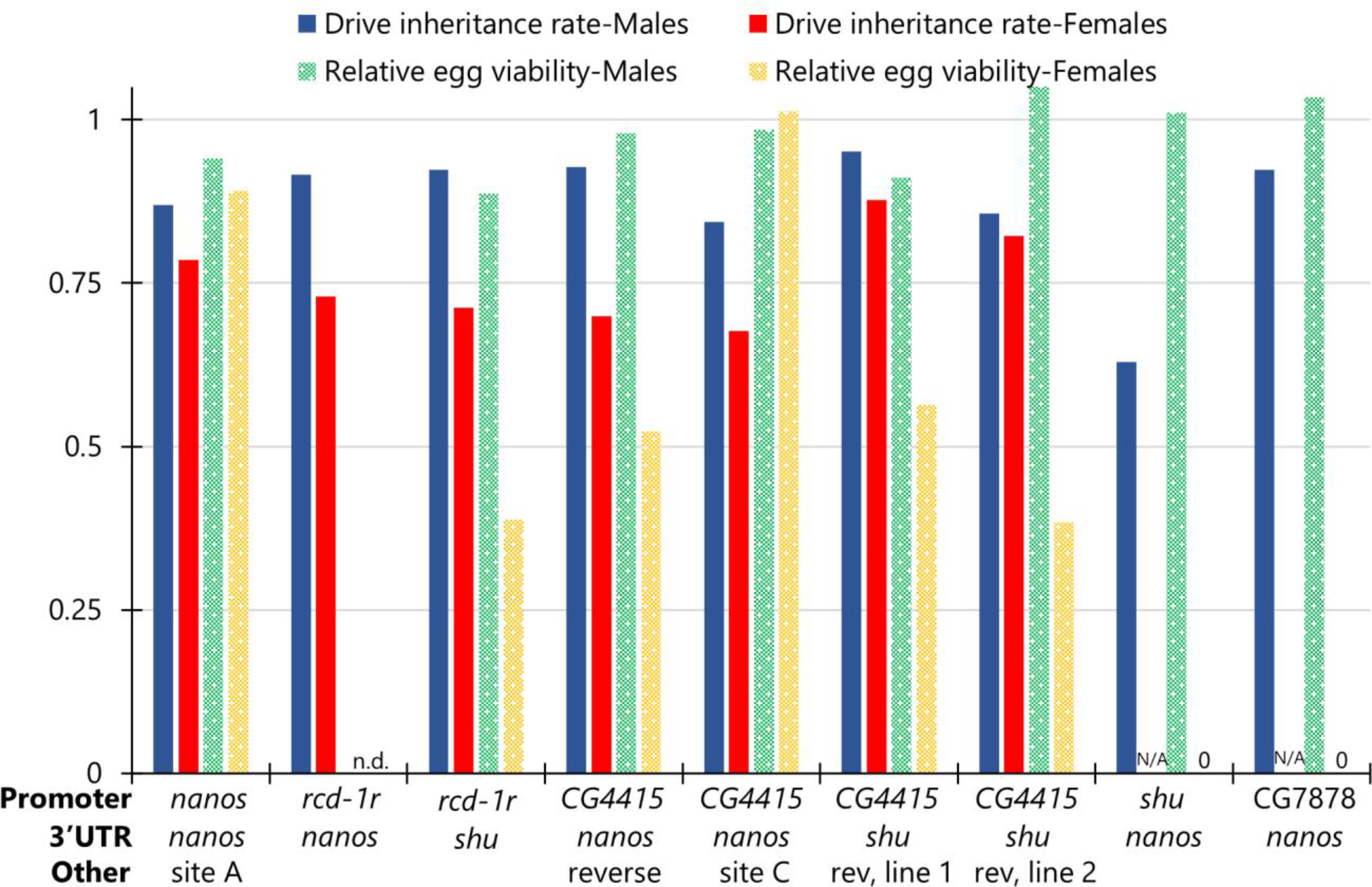
Drive inheritance and offspring viability of promoters with homing suppression drive. Flies heterozygous for different Cas9 alleles on chromosome 2R (2L for “site C”) and a drive targeting *yellow-G* gene on chromosome 3 were crossed with *w*^1118^ flies. *yellow-G* is a haplosufficient gene essential for female fertility, so progeny from females suffering from high somatic Cas9 expression (or potentially high germline expression) have lower viability. Progeny were phenotyped for DsRed (drive), and for several experiments, eggs were counted after one day of egg laying. Relative egg viability is the relative rates of egg survival compared to control experiments in which egg viability was measured for drive heterozygotes without Cas9, Cas9 heterozygotes, and crosses between *w*^1118^ flies. “Reverse” indicates that the orientation on one gene of the allele is reversed so that the Cas9 promoter and 3xP3 of EGFP are not adjacent. n.d. - not determined, N/A - not applicable (no offspring to measure inheritance), 0 - no viable eggs. The leftmost drive data is from a previous study^14^.

For two drives, embryo resistance as inferred by measuring if female drive carrier progeny of females with the drive and Cas9 were fertile. Infertility would be caused if the wild-type copy of *yellow-G* that these progeny receive from their father is converted into a nonfunctional resistance allele in the early embryo. This could be detected by allowing the fly to mate with males and lay eggs for a week. If larvae were observed in the vial, then the mother was fertile. For the Cas9 with the *CG4415* promoter, *nanos* 3’ UTR, and reverse orientation, all 36 females tested were fertile (as well as 21 female controls that were similar, but the offspring of males with drive and Cas9). When the *rcd-1r* promoter was used with the *shu* 3’ UTR, all 13 females were fertile (together with 23 control females, as above). This indicates that the suppression drive targeting *yellow-G* likely has naturally lower embryo resistance than the drives targeting EGFP or *yellow*, allowing our promoters with lower embryo resistance in these other systems to also avoid high embryo resistance with the suppression drive.

### Suppression drive cage experiments

Previously, our 4-gRNA homing suppression drive targeting *yellow-G* failed to spread in two cage populations, reaching only an intermediate equilibrium frequency^14^. The *nanos* promoter only showed small fitness costs in individual crosses, but it had substantially higher fitness costs in cage populations, perhaps due to different environmental conditions such as increased desiccation risk. High embryo resistance in the *nanos* promoter also contributed to poor performance. We selected two split Cas9 lines at nearly the same genomic site for similar cage experiments, one with the *CG4415* promoter and *nanos* 3’ UTR (with Cas9 in the same orientation as EGFP), and another with the *rcd-1r* promoter and *shu* 3’ UTR (line #1 for higher performance).

First, Cas9 homozygous female virgins were collected, mixed, and then were mated to either drive heterozygous males or males that were wild-type at the drive site (all males were also homozygous for Cas9). Then, males were removed, and females were allowed to lay eggs in cage bottles for two days. Females were removed, and new food was provided to offspring eleven days later, and these offspring were considered to be “generation zero,” in which the drive heterozygote frequency was approximately 10%. Flies were then kept on a 12-day cycle with discrete generations and approximately 24 hours of egg laying per generation. All flies were phenotyped to track the drive carrier frequency and total population size.

In the cage with Cas9 driven by the *rcd-1r* promoter, drive carrier frequency slowly increased but always remained lower than 27%, apparently reaching a low equilibrium value (Figure 7, Data Set S6). The total population size was not affected, fluctuating from a maximum of 3019 adults to a minimum of 840. However, for the other cage with Cas9 element driven by *CG4415* promoter (Figure 7), the drive carrier frequency increased quickly, then remained constant for a few generations (possibly due to random fluctuations or differences in food characteristics; environmental fluctuations are unlikely because flies were maintained in a temperature and humidity-controlled environment). Finally, the drive frequency continued to increase, and in this last phase after generation 9, the population was steadily reduced, falling to zero in generation 14. This result was likely due to reduced embryo resistance, but fitness costs were also perhaps different than in individual cross experiments.

**Figure 7.**
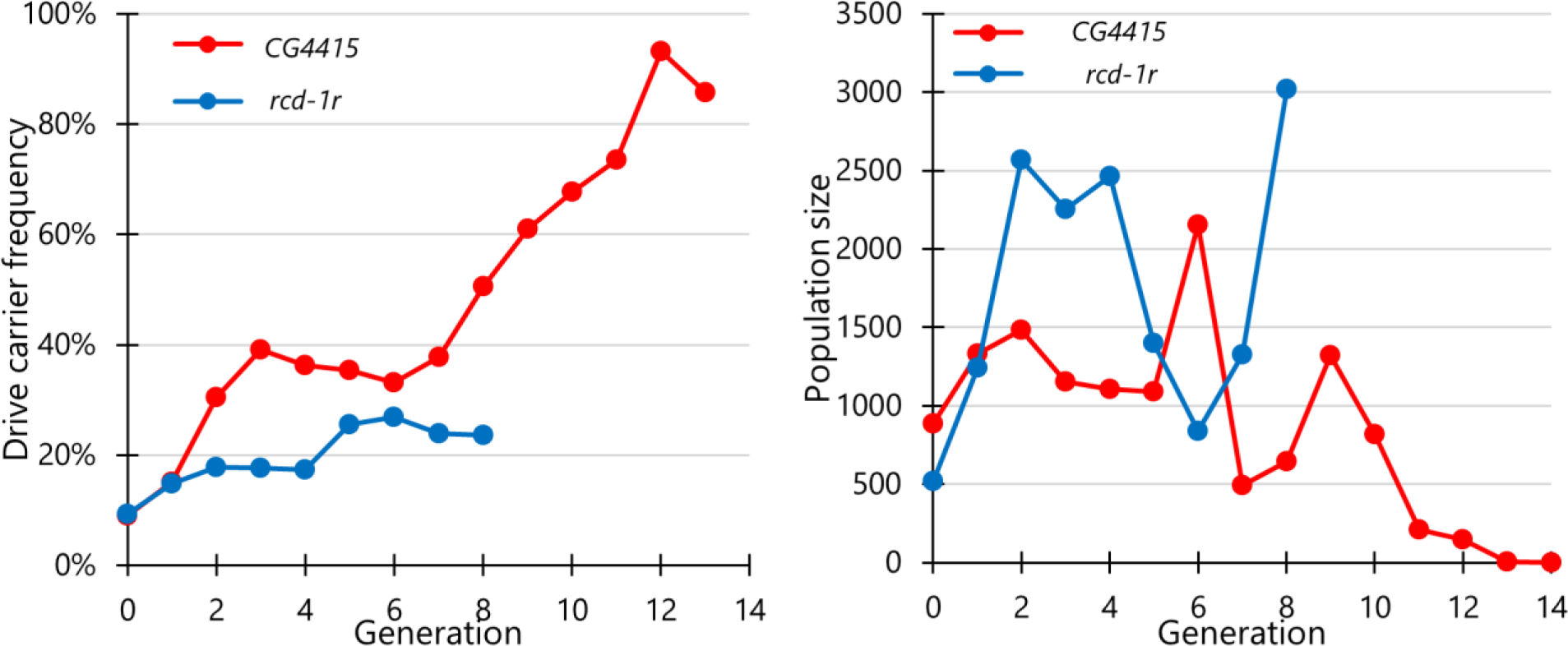
Multigenerational cage experiments with homing suppression drive. Cage experiments were initialized in generation zero with approximately 10% of flies carrying one copy of the drive allele and 90% flies that lacked a drive allele. All individuals were homozygous for the Cas9 allele (either *CG4415* with the *nanos* 3’ UTR in reverse orientation, or *rcd-1r* line #1 with the *shu* 3’ UTR). The cage populations were maintained separately with nonoverlapping generations, each lasting 12-13 days with 1 day for egg laying. All individuals for each generation were phenotyped for DsRed (indicating drive carriers that could be homozygous or heterozygous), and the total population was also recorded.

To assess drive performance parameters based on cage data, a maximum likelihood method was applied, similarly to previous studies^14, 15, 24, 39^. We used a simple model with one gRNA and no functional resistance, the latter of which is likely a valid assumption due to four gRNAs^14^. Assuming only one gRNA may slightly underestimate drive performance^9^. Females and males were assumed to have a different drive conversion efficiency based on drive inheritance data (Dats Set S5), though performance in cage Cas9 homozygotes may be slightly different than in individuals crosses where flies only had one copy of Cas9. Embryo resistance was set at 5% for both drives. Varying this parameter between 0% and 10% had little effect on the results. The fitness of female drive heterozygous was allowed to vary and was inferred by the model.

With a fixed effective population size, the model inferred a low effective population size for the *CG4415* cage, indicating a poor fit (Table S3). This was notably improved if the last two generations were removed, and further improved if the transitions between generations 3 and 6 were removed (when the drive frequency slightly declined instead of increasing as expected). We reasoned that the large population size changes in this suppression drive, especially near the end, could be negatively affecting a model with fixed effective population size. We therefore allowed it to vary, assuming that it was a fixed percentage of the average between the two generations for each generation transition. This produced a better model that was not substantially improved by removing the last two generations, and only slightly improved by removing the middle generations. With all generations, the effective population size was 4.4% of the census size, and the female fitness was estimated as 0.7 (95% confidence interval: 0.38 to 1.14). With the unusual middle transition removed, the effective population size was 5.3%, and the female heterozygote fitness was 0.96 (95% confidence interval: 0.53 to 1.58). For the *rcd-1r* cage, the model was a better fit, with a higher effective population size and an inferred fitness of 0.3 (with a 95% confidence interval that ended well below 1) (Table S3). It is quite possible that fitness changed substantially with the state of the food during the experiment (despite using the same recipe), with time, and certainly between this study and a previous study^14^, which used a similar recipe but different sources of ingredients. Nevertheless, the fitness cost in the *CG4415* cage appeared to be substantially less than *nanos* or *rcd-1r*, and coupled with the greatly reduced rate of embryo resistance compared to *nanos* (which was likely over 50% for this target site^14^), this was sufficient to allow suppression of a large, robust, *Drosophila* cage population.

## Discussion

In this study, we compared several Cas9 promoters in CRISPR homing gene drives. The previously well-characterized *nanos* promoter had high germline cut rates and undetectable somatic expression but suffered from high embryo resistance allele formation due to maternally deposited Cas9. Among the eleven other promoters we tested, *rcd-1r*, *CG4415*, *shu*, and *mei-W68* could often achieve similar drive inheritance rates but substantially lower embryo resistance rates compared to *nanos*. However, *shu* had trouble maintaining high drive inheritance in systems with less active gRNAs, and *mei-W68* still had moderate embryo resistance. All of these suffered from somatic expression, but this was limited enough in *CG4415* that it was suitable for use in a suppression drive, resulting in successful population elimination.

To assess drive performance, visual markers allow genotyping based on phenotypes, avoiding laborious sequencing. Our EGFP target site drives and *yellow* split drives are both suitable for this purpose, and each has its advantages. Neither affects fly viability, but only EGFP allows assessment of male drive performance. Split Cas9 lines can be used more flexibly with other drive lines, and *yellow* allows easier assessment of female germline performance and somatic expression. It is also potentially more representative of somatic expression in a wider array of tissues. However, we saw similar somatic patterns in both of these drives. While our promoter choices were chosen for their strong germline expression and low expression in other tissues, they still have some non-germline expression^45^, which can be in somatic cells or lead to embryo resistance. It is unclear if this native expression level would, in fact, be enough to produce the cleavage patterns we observed, or if our Cas9 elements had different expression patterns due to missing regulatory sequences or other factors.

Genomic location certainly has a substantial effect on expression, as indicated by our tests with an alternate genomic site for the *CG4415* promoter split Cas9 element, which had lower somatic expression (though germline performance was also slightly worse, so it is unclear if tissue-specific expression was changed or if expression was just generally lower). Indeed, while we generally desire high germline cut rates, there may be considerable incentive to cease further increases in Cas9 expression once this has been achieved (in other words, aim for the minimum level of Cas9 expression needed to achieve high drive conversion rates). Insertion of PEST sequence at the C-terminus of Cas9 decreased embryo resistance rates somewhat in the EGFP target sites drive without affecting germline performance. However, when Cas9 expression was lower in the *yellow* system with other promoters (direct measurement of germline cutting usually remained near 100%, but we infer this from lower baseline embryo resistance rates from maternally deposited Cas9 that was originally expressed in the germline), addition of a PEST sequence severely reduced Cas9 germline drive conversion activity. This is potentially also supported by our observation in this study and previous ones^6, 14, 15, 19^ that drive conversion was usually higher in males than in females for the *RpL35A* and *yellow-G* drives, but usually higher in females than males in EGFP target drives or a drive targeting *cinnabar*. Embryo resistance, on the other hand, was higher in the EGFP and *cinnabar* drives. If embryo resistance is closely correlated with germline expression, then it is possible that generally higher Cas9 expression in males can explain these results. When expression is low, embryo resistance is low, and females may not have high germline cutting, leading to persistence of many wild-type alleles and reduced drive conversion. Males with higher expression may still achieve higher drive conversion rates. However, as expression increases in both sexes, females now have higher drive conversion, while male drive conversion is actually reduced. With males now having more than sufficient germline expression, cleavage would tend to occur earlier on average. Resistance alleles are known to form in pre-gonial germline cells, and cleavage at this temporal phase may tend to produce more resistance alleles compared to drive conversion than later cleavage in or closer to the gametocyte stage. However, additional data would be needed for such a hypothesis to be strongly supported.

We also assessed alternate 3’ UTRs, albeit less systematically than promoters. It is possible that the 3’ UTR may interact with other regulatory elements at the mRNA stage, and it could also influence the rate of mRNA degradation. However, we generally found that the *nanos* 3’ UTR reduces embryo resistance and somatic expression compared to 3’ UTR elements matching the promoter, at least for a few of our promoters that already had low embryo resistance.

Haplolethal rescue drives have substantial advantages over ones targeting a haplosufficient gene. Resistance alleles are more easily eliminated in haplolethal drive system^15^, and the drive can reach 100% final frequency even if it has fitness costs (if fitness costs are present, the drive carrier frequency will be 100%, but total drive allele frequency will be less, similarly to CRISPR toxin-antidote drives^38, 46, 47^). However, embryo resistance can remove drive alleles, and somatic expression can form nonfunctional resistance alleles, leading to nonviability or heavy fitness costs (Table 1). In our haplolethal drive system, despite lower embryo cut rates than in other systems^15^, promoters with high embryo resistance prevented successful egg production by females. However, we found several promoters with lower embryo resistance that appeared to also have no detectable negative effects from somatic expression. While somatic expression can certainly lead to heavy fitness costs in haplolethal drive systems^48^, in this case, we tested them in homing drive systems, where drive conversion is possible in somatic cells^5^. Drive conversion would provide a second copy of the rescue gene, resulting in healthy cells. This, combined with the naturally low cut rates of this drive, likely allowed them to avoid detectable fitness costs, which is quite promising for future use of haplolethal homing drives in other species.

**Table 1.** Most impactful consequences of imperfect drive performance.

| **Problem** | **Haplolethal modification** | **Haplosufficient modification** | **Female-fertility Suppression** |
| --- | --- | --- | --- |
| *low drive conversion* | slower drive | slower drive | much lower power |
| *high germline resistance* | faster drive | faster drive | slower drive |
| *high embryo resistance* | possible failure | slower drive | lower power* |
| *high somatic expression* | fitness cost<br>possible failure | possible small fitness cost | large fitness cost<br>lower power* |

\*only if drive conversion is not very high

Though less problematic than functional resistance alleles, nonfunctional resistance alleles are more difficult to address and a primary obstacle for creating good gene drive systems, especially in suppression drives. All homing suppression drives targeting female fertility thus far have suffered from fitness costs in drive heterozygous females. This is critically important^49^ because it reduces the genetic load (suppression power) of the drive when drive conversion in not nearly 100%^50^, which can result in persistence of the population (Table 1). Even with high drive conversion, it can complicate suppression in spatial environments^44^. Embryo resistance has a similar effect, sterilizing daughters of female drive heterozygotes. There is thus a higher incentive to develop improved promoters for suppression drives than otherwise well-designed modification drives. Previously, our 4-gRNA drive targeting *yellow-G* failed to suppress a cage when paired with the *nanos* promoter^14^, but with the *CG4415* promoter driving Cas9, suppression was successful. The main advantage of *CG4415* was the far lower rate of embryo resistance, but it appeared to have less fitness costs in heterozygotes in the cage study than *nanos* as well, unlike our test with *rcd-1r*, which performed worse than *nanos* despite also reducing embryo resistance. This is also somewhat contradictory to our egg viability experiments, though the 95% confidence intervals of fitness overlap, and different conditions in the cage experiments may result in different actual fitness costs compared to individual crosses, particularly since *yellow-G* is needed for egg shells, meaning that environmental conditions may strongly affect fitness. The observation of lower fitness costs with *CG4415* compared to *nanos* is also unexpected considering the lower somatic expression in *nanos*. However*, yellow-G* is expressed in the ovaries, even if perhaps not in gametes. It is possible that even though *nanos* has lower general somatic expression, *yellow-G* was still disrupted in some ovary cells where it was needed, while the lower germline expression of *CG4415* (again, based on embryo resistance) caused less disruption to these cells and thus less fitness cost, allowing high efficiency and rapid success in the cage population.

While performance of our promoters has revealed useful general information in the model organism *D. melanogaster*, they could potentially be applied to other species as well. This certainly seems to be the case with U6 promoters, which have been used to express gRNAs in every CRISPR gene drive study thus far. However, these have a less complex required expression pattern than Cas9, where we often desire restriction of cleavage activity to the germline, rather than just accepting high expression everywhere. Recently, a drive system targeting *doublesex* was tested in the major crop pest *D. suzukii* and yielded good results with the *nanos* promoter for Cas9^51^. Performance was actually better than in *D. melanogaster* for drive conversion, though this could have been caused by addition of a second nuclear localization signal. It is possible that other promoters would have similar performance in this and other closely related species, which could include important pests such as the medfly. However, in more distantly related species such as mosquitoes, the situation is different. In an *Anopheles* homing suppression drive^10^, the *zpg* promoter lacked the high embryo resistance and somatic expression that we saw in our studies. *nanos* also had higher drive conversion than in *D. melanogaster* and had much lower embryo resistance^30^. *vasa* had high embryo resistance and somatic expression in both species^29^. The *shu* promoter in *Aedes aegypti* could support very high cut rates and drive conversion in the germline in some lines, while most other promoters failed to achieve this^17, 27, 28^. This contrasts with our results, where *shu* was a weaker promoter, achieving high efficiency only in the EGFP target line. All these comparisons, and our promoter assessment in general, have the important caveat of the exact length of promoter elements utilized. In some of our promoters in particular, we sought to use shorter elements to avoid the coding sequence of other genes in an attempt to find more compact regulatory elements and avoid fitness costs from undesired transcription in different directions.

Overall, our study demonstrated that homing drives could achieve high efficiency with several germline promoters, 5’ UTRs, and 3’UTR regulatory elements for Cas9. In multiple systems, we identified strengths and weaknesses of new promoters and how they interact with varying drive elements. These regulatory elements could offer large advantages for drive systems in *Drosophila*, and their homologs could even be useful in other species, where they may have substantially varying performance but could still serve as useful candidates.

## Acknowledgements

This study was supported by laboratory startup funds from Peking University, the Center for Life Sciences, the NSFC Overseas Youth Fund, the SLS-Qidong Innovation Fund, and the National Institutes of Health award F32AI138476 to JC.

## Supplemental Information

**Figure S1.**
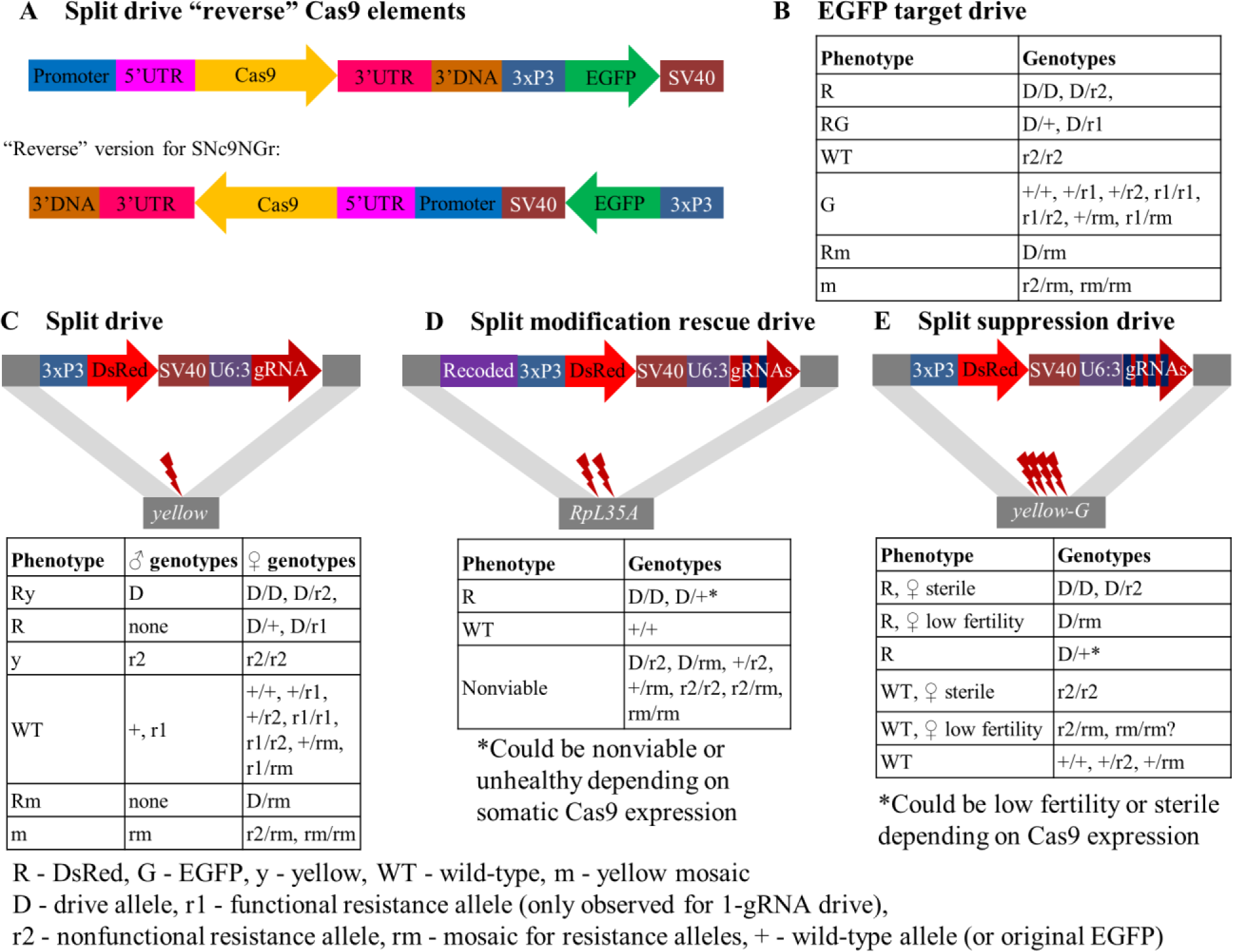
Schematic diagram of additional constructs with rearranged elements. **(A)** In some split Cas9 lines, the orientation of the Cas9 gene is reversed (on in one case, the orientation of the EGFP gene) to prevent interference between the promoter elements of Cas9 and EGFP. (**B**) Drives targeting EGFP are marked with DsRed. The drive and nonfunctional resistance alleles (r2) can disrupt EGFP. Resistance alleles can also be mosaic. Functional resistance alleles (r1) are rare. (**C**) The *yellow* gene has a recessive knockout phenotype that produces a yellow color on the body and wings. Both the drive and nonfunctional resistance alleles can produce this knockout phenotype. The drive also carries a DsRed gene and is designed to be used with a split Cas9 line. Because *yellow* is on the X-chromosome, males have simpler genotypes, while female phenotypes are more complex and influenced by factors such as leaky somatic Cas9 expression and maternal Cas9 deposition. This drive produces perhaps ∼10% functional resistance alleles. (**D**) The split rescue drive targets the haplolethal gene *RpL35A*. Flies with nonfunctional resistance alleles are nonviable. Somatic Cas9 expression can result in resistance allele formation or drive conversion, so higher amounts are needed to reduce viability. (**E**) The split suppression drive targets and disrupts yellow-G, a haplosufficient female fertility gene. Thus, only females with wild-type alleles are fertile. Somatic Cas9 expression substantially reduces fertility.

**Figure S2.**
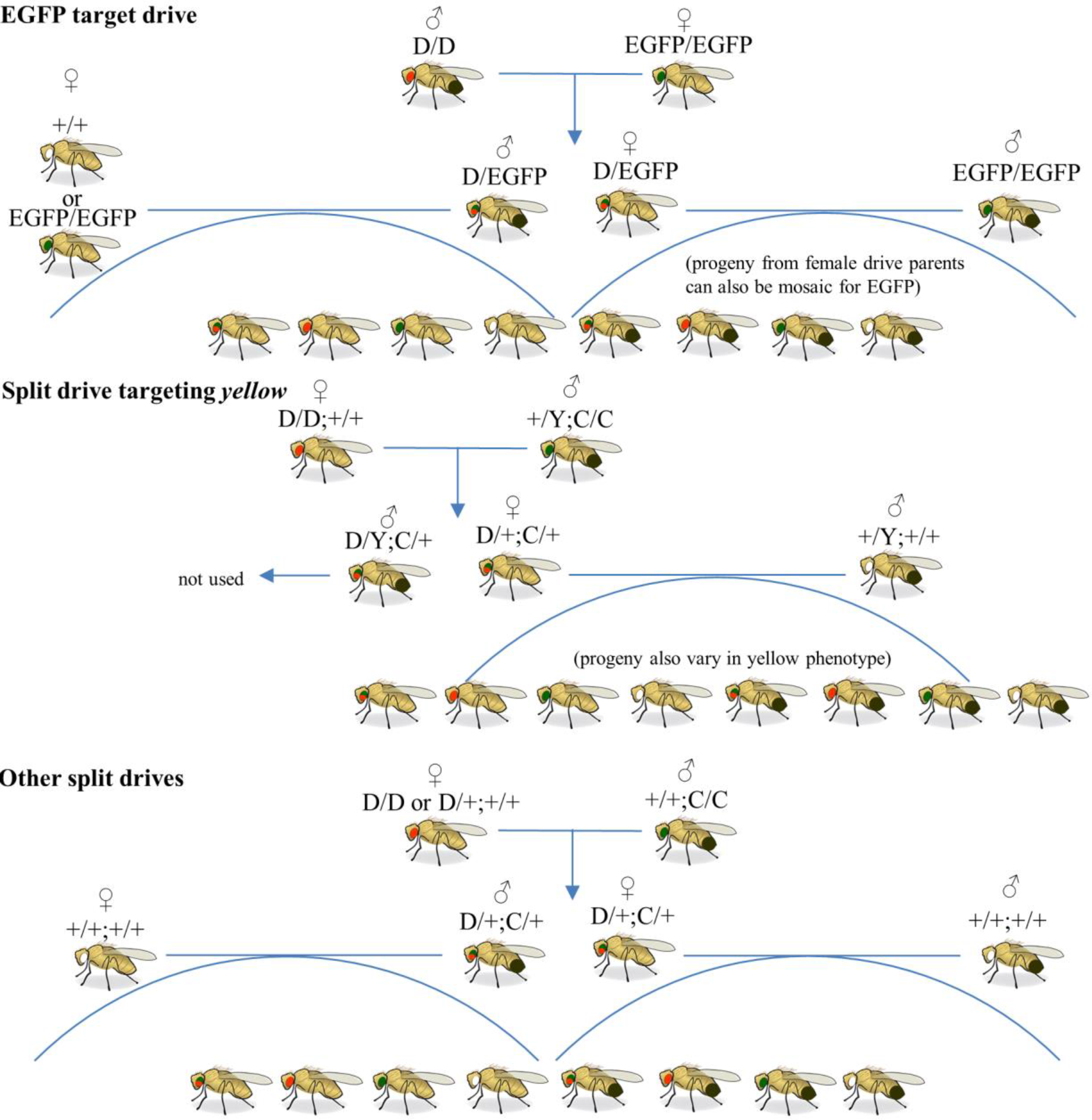
Crossing scheme. The figure shows the crossing scheme for each class of drive in the study. Because maternal Cas9 and gRNA generate embryo resistance alleles, only drive males were used to start the crosses for the EGFP target drive so that drive heterozygous offspring would all be drive/wild-type (instead of potentially drive/resistance). The split drives were not affected by this issue because gRNA is also required to be maternally deposited for embryo resistance alleles to form. For the split drive targeting *yellow*, only female heterozygotes were assessed for drive performance because the drive is X-linked.

**Table S1.** List of donor plasmid names with Cas9 regulatory features.

SN = split Cas9 line, BHD = drive targeting EGFP, \*except for BHDaaN, which is split Cas9 at site A
| **Line/Donor Plasmid** | **Promoter/5' UTR** | **3' UTR/Terminator** | **Other Feature** |
| --- | --- | --- | --- |
| SNc9VnG | *vasa* | *nanos* |  |
| SNc9VnGp | *vasa* | *nanos* | PEST |
| SNc9NG | *nanos* | *nanos* |  |
| SNc9NGr | *nanos* | *nanos* | reverse |
| SNc9NsG | *nanos* | *shu* |  |
| SNc9NvG | *nanos* | *vasa* |  |
| BHDgN1cv3 | *nanos* | *nanos* |  |
| BHDgN1p | *nanos* | *nanos* | PEST |
| BHDaaN* | *nanos* | *nanos* | *split Cas9 line |
| SNc9DnG | *rcd-1r* | *nanos* |  |
| SNc9DG | *rcd-1r* | *rcd-1r* |  |
| SNc9DsG | *rcd-1r* | *shu* |  |
| SNc9DpG | *rcd-1r* | *nanos* | PEST |
| SNc9DpGr | *rcd-1r* | *nanos* | reverse, PEST |
| BHDgD1 | *rcd-1r* | *nanos* |  |
| BHDgD1d | *rcd-1r* | *rcd-1r* |  |
| SNc9XnGr | *CG4415* | *nanos* |  |
| SNc9XG | *CG4415* | *CG4415* |  |
| SNc9XSGr1 | *CG4415* | *shu* | reverse, line 1 |
| SNc9XSGr2 | *CG4415* | *shu* | reverse, line 2 |
| SNc9XGd | *CG4415* | *rcd-1r* |  |
| SNc9XpG | *CG4415* | *nanos* | reverse, PEST |
| SNc9XpGv2 | *CG4415* | *nanos* | PEST |
| SNcc9XpG | *CG4415* | *nanos* | PEST, site C |
| SNcc9XnG | *CG4415* | *nanos* | site C |
| BHDgX1 | *CG4415* | *nanos* |  |
| BHDgX1x | *CG4415* | *CG4415* |  |
| SNc9ZG | *zpg* | *zpg* |  |
| BHDgZ1 | *zpg* | *nanos* |  |
| BHDgZ1z | *zpg* | *zpg* |  |
| SNc9SnG | *shu* | *nanos* |  |
| SNc9SpG | *shu* | *nanos* | reverse, PEST |
| SNc9SpGv2 | *shu* | *nanos* | PEST |
| BHDgS1 | *shu* | *nanos* |  |
| BHDgS1s | *shu* | *shu* |  |
| SNc9FnG | *CG17658* | *nanos* |  |
| SNc9FpG | *CG17658* | *nanos* | PEST |
| SNc9CnG | *CG7878* | *nanos* |  |
| SNc9CpG | *CG7878* | *nanos* | PEST |
| SNc9EnG | *CG3223* | *nanos* |  |
| BHDgB1 | *β2-tubulin* | *nanos* |  |
| BHDgM1 | *mei-W68* | *nanos* |  |

**Table S2.**
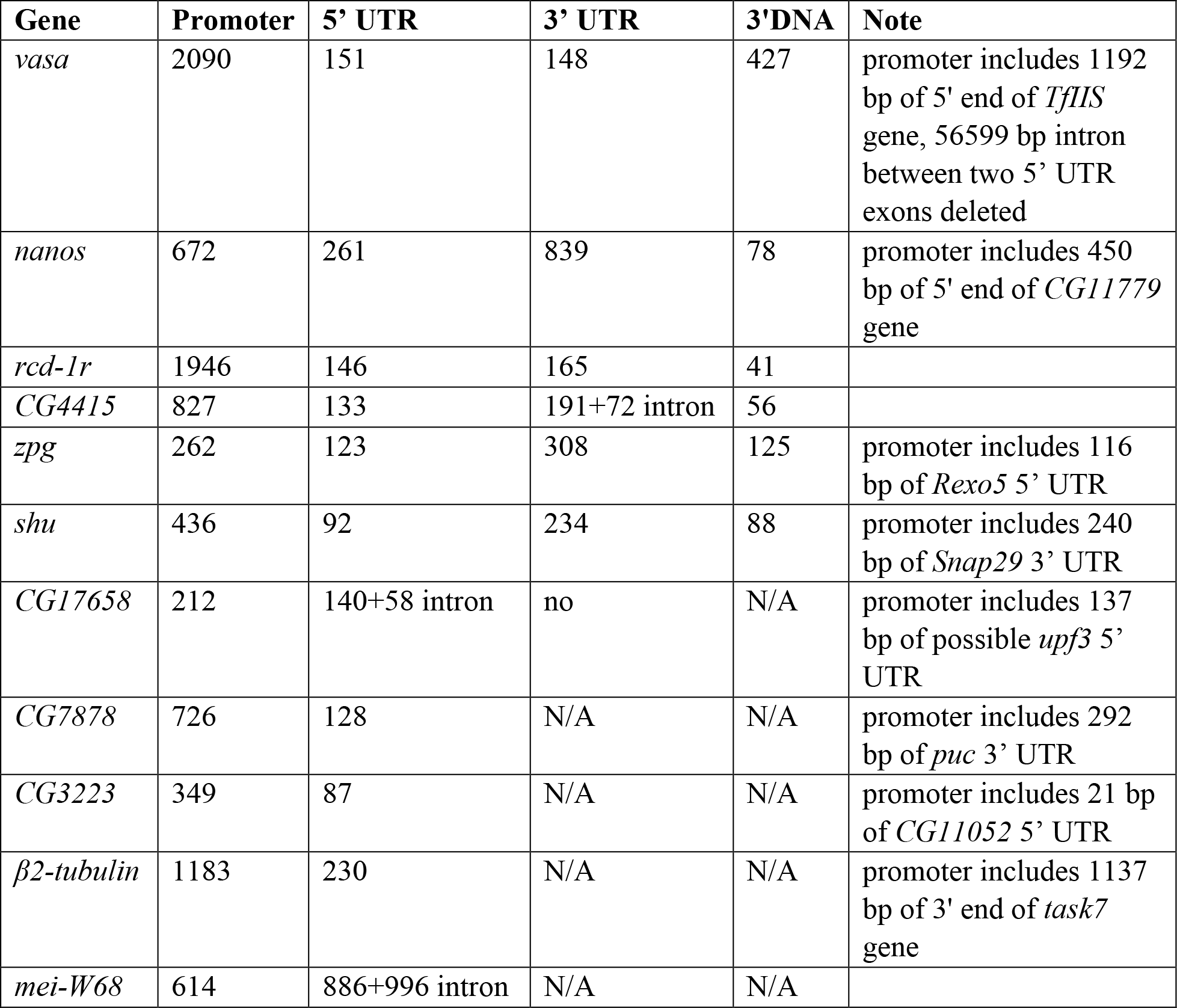
Sizes of Cas9 regulatory elements.

**Figure S3.**
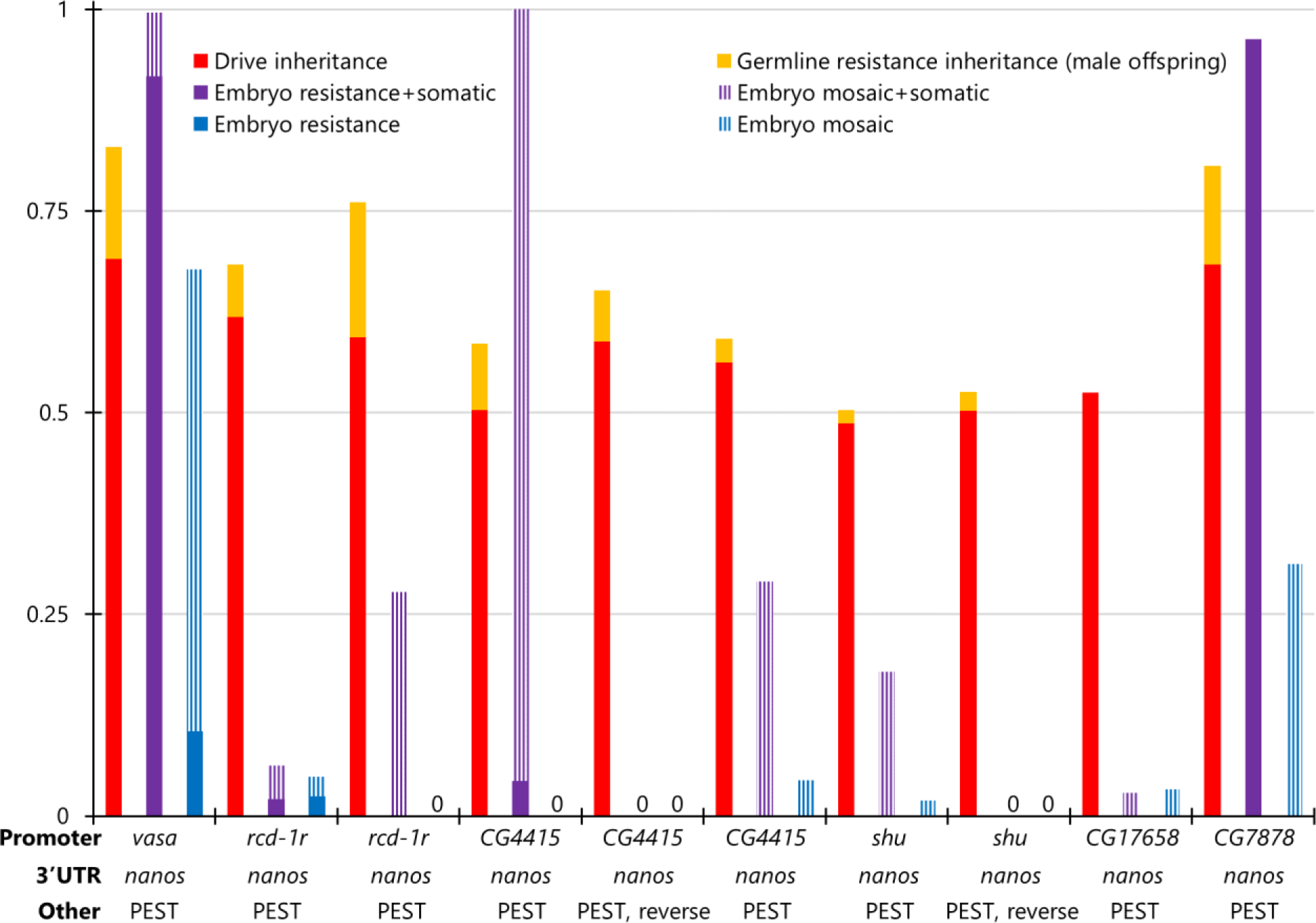
*yellow* target site drive performance with PEST domain. Females heterozygous for different Cas9 alleles on chromosome 2R and heterozygous for a drive targeting the X-linked *yellow* gene were crossed with *w*^1118^ males. Their progeny were phenotyped for DsRed (drive), EGFP (Cas9), and yellow body color. The germline resistance inheritance rate shows the fraction of male progeny that had yellow body color but no drive allele (such flies could also form from embryo resistance allele formation). The fraction of offspring with yellow phenotype (or with mosaic phenotype), inheriting the drive, and also inheriting Cas9 is labeled as “Embryo resistance+somatic” because either maternally deposited Cas9 /gRNA or somatic expression could be responsible for the yellow phenotype. “Embryo resistance rate” (and the corresponding mosaic rate) is similar, but reports the fraction of drive offspring lacking Cas9 that have the yellow phenotype, which can only be caused by maternal deposition. “Reverse” indicates that the orientation on one gene of the allele is reversed so that the Cas9 promoter and 3xP3 of EGFP are not adjacent. 0 - no apparent phenotype in any offspring from embryo resistance or from combined embryo resistance and somatic expression.

**Table S3.** Maximum likelihood estimates of female drive fitness.

| **Model** | **Log-Likelihood** | **AICc** | **Effective Population** | **Female D/+ Fitness** |
| --- | --- | --- | --- | --- |
| Normal | 14.2 | -23.2 | 29<br>[12, 58] | 0.84<br>[0.41, 1.49] |
| Generations 12-13 removed | 14.6 | -23.7 | 53<br>[20, 110] | 0.73<br>[0.41, 1.16] |
| Generations 4-5 removed | 11.6 | -17.9 | 31<br>[11, 67] | 1.00<br>[0.55, 2.21] |
| Gen 4-5, 12-13 removed | 12.2 | -18.9 | 76<br>[24, 177] | 1.00<br>[0.61, 1.57] |
| Normal | 15.3 | -25.4 | 4.4%<br>[1.5%, 8.8%] | 0.70<br>[0.38, 1.14] |
| Generations 12-13 removed | 13.8 | -22.1 | 4.4%<br>[1.5%, 9.3%] | 0.67<br>[0.36, 1.11] |
| Generations 4-5 removed | 12.4 | -19.6 | 5.3%<br>[1.8%, 11.5%] | 0.96<br>[0.53, 1.58] |

***rcd-1r* Cage**
| **Model** | **Log-Likelihood** | **AICc** | **Effective Population** | **Female D/+ Fitness** |
| --- | --- | --- | --- | --- |
| Normal | 15.2 | -24.0 | 126<br>[39, 291] | 0.31<br>[0.09, 0.58] |
| Normal | 15.5 | -24.5 | 8.3%<br>[2.4%, 19.2%] | 0.29<br>[0.09, 0.54] |

AICc - Akaike information criterion, corrected
Brackets show 95% confidence intervals

## References

1. Champer J, Buchman A, Akbari OS. Cheating evolution: engineering gene drives to manipulate the fate of wild populations. Nat Rev Genet, 17, 146–159, 2016.

2. Wang G-H, Du J, Chu CY, Madhav M, Hughes GL, Champer J. Symbionts and gene drive: two strategies to combat vector-borne disease. Trends Genet, 37, 708–723, 2022.

3. Bier E. Gene drives gaining speed. Nat Rev Genet, 1–18, 2021.

4. Hay BA, Oberhofer G, Guo M. Engineering the composition and fate of wild populations with gene drive. Annu Rev Entomol, 66, 407–434, 2021.

5. Li Z, Marcel N, Devkota S, Auradkar A, Hedrick SM, Gantz VM, Bier E. CopyCatchers are versatile active genetic elements that detect and quantify inter-homolog somatic gene conversion. Nat Commun, 12, 1–12, 2021.

6. Champer J, Liu J, Oh SY, Reeves R, Luthra A, Oakes N, Clark AG, Messer PW. Reducing resistance allele formation in CRISPR gene drive. Proc Natl Acad Sci, 115, 5522–5527, 2018.

7. Champer J, Reeves R, Oh SY, Liu C, Liu J, Clark AG, Messer PW. Novel CRISPR/Cas9 gene drive constructs reveal insights into mechanisms of resistance allele formation and drive efficiency in genetically diverse populations. PLoS Genet, 13, e1006796, 2017.

8. Gantz VM, Jasinskiene N, Tatarenkova O, Fazekas A, Macias VM, Bier E, James AA. Highly efficient Cas9-mediated gene drive for population modification of the malaria vector mosquito Anopheles stephensi. Proc Natl Acad Sci U S A, 112, E6736–E6743, 2015.

9. Champer SE, Oh SY, Liu C, Wen Z, Clark AG, Messer PW, Champer J. Computational and experimental performance of CRISPR homing gene drive strategies with multiplexed gRNAs. Sci Adv, 6, eaaz0525, 2020.

10. Kyrou K, Hammond AM, Galizi R, Kranjc N, Burt A, Beaghton AK, Nolan T, Crisanti A. A CRISPR-Cas9 gene drive targeting doublesex causes complete population suppression in caged Anopheles gambiae mosquitoes. Nat Biotechnol, 36, 1062–1066, 2018.

11. Shapiro RS, Chavez A, Porter CBM, Hamblin M, Kaas CS, DiCarlo JE, Zeng G, Xu X, Revtovich A V., Kirienko N V., Wang Y, Church GM, Collins JJ. A CRISPR–Cas9-based gene drive platform for genetic interaction analysis in Candida albicans. Nat Microbiol, 3, 73–82, 2018.

12. Yan Y, Finnigan GC. Analysis of CRISPR gene drive design in budding yeast. *Access Microbiol*, acmi000059, 2019.

13. Weitzel AJ, Grunwald HA, Ceri W, Levina R, Gantz VM, Hedrick SM, Bier E, Cooper KL. Meiotic Cas9 expression mediates gene conversion in the male and female mouse germline. PLOS Biol, 19, e3001478, 2021.

14. Yang E, Metzloff M, Langmüller AM, Xu X, Clark AG, Messer PW, Champer J. A homing suppression gene drive with multiplexed gRNAs maintains high drive conversion efficiency and avoids functional resistance alleles. G3 Genes|Genomes|Genetics, 2022.

15. Champer J, Yang E, Lee E, Liu J, Clark AG, Messer PW. A CRISPR homing gene drive targeting a haplolethal gene removes resistance alleles and successfully spreads through a cage population. Proc Natl Acad Sci, 117, 24377–24383, 2020.

16. Adolfi A, Gantz VM, Jasinskiene N, Lee HF, Hwang K, Terradas G, Bulger EA, Ramaiah A, Bennett JB, Emerson JJ, Marshall JM, Bier E, James AA. Efficient population modification gene-drive rescue system in the malaria mosquito Anopheles stephensi. Nat Commun, 11, 1–13, 2020.

17. Anderson MAE, Gonzalez E, Ang JXD, Shackleford L, Nevard K, Verkuijl SAN, Edgington MP, Harvey-Samuel T, Alphey L. Closing the gap to effective gene drive in Aedes aegypti by exploiting germline regulatory elements. Nat Commun, 14, 1–9, 2023.

18. Hammond AM, Galizi R. Gene drives to fight malaria: current state and future directions. Pathog Glob Health, 111, 412–423, 2017.

19. Champer J, Chung J, Lee YL, Liu C, Yang E, Wen Z, Clark AG, Messer PW. Molecular safeguarding of CRISPR gene drive experiments. Elife, 8, 2019.

20. Noble C, Min J, Olejarz J, Buchthal J, Chavez A, Smidler AL, DeBenedictis EA, Church GM, Nowak MA, Esvelt KM. Daisy-chain gene drives for the alteration of local populations. Proc Natl Acad Sci, 201716358, 2019.

21. Verkuijl SAN, Anderson MAE, Alphey L, Bonsall MB. Daisy-chain gene drives: The role of low cut-rate, resistance mutations, and maternal deposition. PLOS Genet, 18, e1010370, 2022.

22. Sudweeks J, Hollingsworth B, Blondel D V, Campbell KJ, Dhole S, Eisemann JD, Edwards O, Godwin J, Howald GR, Oh KP, Piaggio AJ, Prowse TAA, Ross J V, Saah JR, Shiels AB, Thomas PQ, Threadgill DW, Vella MR, Gould F, Lloyd AL. Locally fixed alleles: A method to localize gene drive to island populations. Sci Rep, 9, 15821, 2019.

23. Dhole S, Lloyd AL, Gould F. Tethered homing gene drives: A new design for spatially restricted population replacement and suppression. Evol Appl, eva.12827, 2019.

24. Metzloff M, Yang E, Dhole S, Clark AG, Messer PW, Champer J. Experimental demonstration of tethered gene drive systems for confined population modification or suppression. BMC Biol, 20, 1–13, 2022.

25. Carballar-Lejarazú R, Ogaugwu C, Tushar T, Kelsey A, Pham TB, Murphy J, Schmidt H, Lee Y, Lanzaro GC, James AA. Next-generation gene drive for population modification of the malaria vector mosquito, Anopheles gambiae. Proc Natl Acad Sci U S A, 117, 22805–22814, 2020.

26. Li M, Yang T, Kandul NP, Bui M, Gamez S, Raban R, Bennett J, Sánchez C HM, Lanzaro GC, Schmidt H, Lee Y, Marshall JM, Akbari OS. Development of a confinable gene drive system in the human disease vector Aedes aegypti. Elife, 9, e51701, 2020.

27. Verkuijl SAN, Gonzalez E, Li M, Ang JXD, Kandul NP, Anderson MAE, Akbari OS, Bonsall MB, Alphey L. A CRISPR endonuclease gene drive reveals distinct mechanisms of inheritance bias. Nat Commun, 13, 1–10, 2022.

28. Reid W, Williams AE, Sanchez-Vargas I, Lin J, Juncu R, Olson KE, Franz AWE. Assessing single-locus CRISPR/Cas9-based gene drive variants in the mosquito Aedes aegypti via single generation crosses and modeling. G3 Genes|Genomes|Genetics, 2022.

29. Hammond A, Galizi R, Kyrou K, Simoni A, Siniscalchi C, Katsanos D, Gribble M, Baker D, Marois E, Russell S, Burt A, Windbichler N, Crisanti A, Nolan T. A CRISPR-Cas9 gene drive system targeting female reproduction in the malaria mosquito vector Anopheles gambiae. Nat Biotechnol, 34, 78–83, 2016.

30. Hammond A, Karlsson X, Morianou I, Kyrou K, Beaghton A, Gribble M, Kranjc N, Galizi R, Burt A, Crisanti A, Nolan T. Regulating the expression of gene drives is key to increasing their invasive potential and the mitigation of resistance. PLOS Genet, 17, e1009321, 2021.

31. Chan YS, Naujoks DA, Huen DS, Russell S. Insect population control by homing endonuclease-based gene drive: an evaluation in Drosophila melanogaster. Genetics, 188, 33–44, 2011.

32. Chan YS, Takeuchi R, Jarjour J, Huen DS, Stoddard BL, Russell S. The design and in vivo evaluation of engineered I-OnuI-based enzymes for HEG gene drive. PLoS One, 8, e74254, 2013.

33. Chan YS, Huen DS, Glauert R, Whiteway E, Russell S. Optimising homing endonuclease gene drive performance in a semi-refractory species: the Drosophila melanogaster experience. PLoS One, 8, e54130, 2013.

34. Simoni A, Siniscalchi C, Chan YS, Huen DS, Russell S, Windbichler N, Crisanti A. Development of synthetic selfish elements based on modular nucleases in Drosophila melanogaster. Nucleic Acids Res, 42, 7461–7472, 2014.

35. Nash A, Capriotti P, Hoermann A, Papathanos PA, Windbichler N. Intronic gRNAs for the construction of minimal gene drive systems. Front Bioeng Biotechnol, 0, 570, 2022.

36. Carrami EM, Eckermann KN, Ahmed HMM, Sánchez C. HM, Dippel S, Marshall JM, Wimmer EA. Consequences of resistance evolution in a Cas9-based sex-conversion suppression gene drive for insect pest management. Proc Natl Acad Sci, 201713825, 2018.

37. Davis MW, Jorgensen EM. ApE, a plasmid editor: A freely available DNA manipulation and visualization program. Front Bioinforma, 0, 5, 2022.

38. Champer J, Lee E, Yang E, Liu C, Clark AG, Messer PW. A toxin-antidote CRISPR gene drive system for regional population modification. Nat Commun, 11, 1082, 2020.

39. Liu J, Champer J, Langmüller AM, Liu C, Chung J, Reeves R, Luthra A, Lee YL, Vaughn AH, Clark AG, Messer PW. Maximum likelihood estimation of fitness components in experimental evolution. Genetics, 211, 1005–1017, 2019.

40. R Core Team. R: A language and environment for statistical computing. www.R-project.org. 2018.

41. Tomancak P, Berman BP, Beaton A, Weiszmann R, Kwan E, Hartenstein V, Celniker SE, Rubin GM. Global analysis of patterns of gene expression during Drosophila embryogenesis. Genome Biol, 8, 1–24, 2007.

42. Galizi R, Doyle LA, Menichelli M, Bernardini F, Deredec A, Burt A, Stoddard BL, Windbichler N, Crisanti A. A synthetic sex ratio distortion system for the control of the human malaria mosquito. Nat Commun, 5, 3977, 2014.

43. Fasulo B, Meccariello A, Morgan M, Borufka C, Papathanos PA, Windbichler N. A fly model establishes distinct mechanisms for synthetic CRISPR/Cas9 sex distorters. PLOS Genet, 16, e1008647, 2020.

44. Champer SE, Kim IK, Clark AG, Messer PW, Champer J. Anopheles homing suppression drive candidates exhibit unexpected performance differences in simulations with spatial structure. Elife, 11, 2022.

45. Brown JB, Boley N, Eisman R, May GE, Stoiber MH, Duff MO, Booth BW, Wen J, Park S, Suzuki AM, Wan KH, Yu C, Zhang D, Carlson JW, Cherbas L, Eads BD, Miller D, Mockaitis K, Roberts J, Davis CA, Frise E, Hammonds AS, Olson S, Shenker S, Sturgill D, Samsonova AA, Weiszmann R, Robinson G, Hernandez J, Andrews J, Bickel PJ, Carninci P, Cherbas P, Gingeras TR, Hoskins RA, Kaufman TC, Lai EC, Oliver B, Perrimon N, Graveley BR, Celniker SE. Diversity and dynamics of the Drosophila transcriptome. Nature, 512, 393–399, 2014.

46. Champer J, Kim IK, Champer SE, Clark AG, Messer PW. Performance analysis of novel toxin-antidote CRISPR gene drive systems. BMC Biol, 18, 27, 2020.

47. Champer J, Champer SE, Kim IK, Clark AG, Messer PW. Design and analysis of CRISPR-based underdominance toxin-antidote gene drives. Evol Appl, eva.13180, 2020.

48. Chen J, Xu X, Champer J. Assessment of distant-site rescue elements for CRISPR toxin-antidote gene drives. Front Bioeng Biotechnol, 11, 1138702, 2023.

49. Wedell N, Price TAR, Lindholm AK. Gene drive: Progress and prospects. Proc R Soc B Biol Sci, 286, 2019.

50. Beaghton AK, Hammond A, Nolan T, Crisanti A, Burt A. Gene drive for population genetic control: non-functional resistance and parental effects. Proceedings Biol Sci R Soc B Biol Sci, 286, 20191586, 2019.

51. Yadav AK, Butler C, Yamamoto A, Patil AA, Lloyd AL, Scott MJ. CRISPR/Cas9-based split homing gene drive targeting doublesex for population suppression of the global fruit pest Drosophila suzukii. Proc Natl Acad Sci, 120, e2301525120, 2023.

